# A non-invasive photoactivatable split-Cre recombinase system for genome engineering in zebrafish

**DOI:** 10.1101/2023.06.23.546268

**Authors:** Ramy Elsaid, Aya Mikdache, Patricia Diabangouaya, Gwendoline Gros, Pedro P. Hernández

**Affiliations:** Institut Curie, PSL Research University CNRS UMR 3215, INSERM U934, 26 Rue d’Ulm, 75248 Paris Cedex 05, France

**Keywords:** Lineage tracing, Zebrafish, Optogenetics, Split-Cre recombinase, Magnets

## Abstract

The cyclic recombinase (Cre)/*loxP* recombination system is a powerful technique for *in vivo* cell labeling and tracking. However, achieving high spatiotemporal precision in cell tracking using this system is challenging due to the requirement for reliable tissue-specific promoters. In contrast, light-inducible systems offer superior regional confinement, tunability and non-invasiveness compared to conventional lineage tracing methods. Here, we took advantage of the unique strengths of the zebrafish to develop an easy-to-use highly efficient, genetically encoded, Magnets-based, light-inducible transgenic Cre/*loxP* system. Our system relies on the reassembly of split Cre fragments driven by the affinity of the Magnets and is controlled by the zebrafish *ubiquitin* promoter. We demonstrate that our system does not exhibit phototoxicity or leakiness in the dark, and it enables efficient and robust Cre/*loxP* recombination in various tissues and cell types at different developmental stages through noninvasive illumination with blue light. Our newly developed tool is expected to open novel opportunities for light-controlled tracking of cell fate and migration *in vivo*.

**Highlights:**

- zPA-Cre is a novel zebrafish transgenic optogenetic split-Cre tool
- zPA-Cre allows loxP-mediated DNA recombination in various tissues and cell types
- It enables light-induced DNA recombination at different developmental stages.
- It allows spatio-temporal DNA recombination in a specific tissue region with high resolution

## Introduction

Cell marking by genetic recombination is a powerful technique for noninvasively tracing cell fate and migration in rapidly evolving contexts such as embryonic development^1,2^. Various recombinases have been used in zebrafish studies^3,4^, however, the Cre/*loxP* system remains the primary method for genetic lineage tracing in this species^2^. The Cre recombinase, derived from the bacteriophage P1, specifically targets two pairs of 34-bp sequences known as *loxP* sites^5^. Due to its simplicity and efficiency, the Cre/*loxP* system has been widely used in applications such as lineage tracing and genome manipulation in transgenic animals^1^. Typically, the Cre recombinase is expressed under a tissue- or cell-specific promoter and can be combined with chemical-inducible systems such as tamoxifen to achieve temporally controlled genetic recombination^1,6,7^. By crossing transgenic lines containing both Cre and *loxP-STOP-loxP* constructs, Cre specifically activates the reporter in tissues or cells expressing the promoter by permanently removing the STOP cassette, thereby enabling the tracing of their progeny^1,2^. Nonetheless, these conventional Cre/*loxP* recombination approaches have some limitations. For instance, it can be challenging to target and trace cells or tissues that lack a specific promoter during development and/or regeneration. Furthermore, the temporal induction of the Cre/*loxP* system using chemicals may have undesirable effects, including cytotoxicity and cellular perturbation^8,9^. Moreover, a critical limitation of conventional Cre/*loxP* recombination systems is their inability to selectively target specific sub-regions of tissues or cells with similar gene expression patterns.

In contrast to conventional lineage tracing approaches, light-inducible systems offer a significant advantage by enabling noninvasive photoactivation in deep tissues with high spatial and temporal resolution^10,11^. Consequently, several attempts have been made to develop optically controlled Cre systems in zebrafish, as reported in previous studies^12–16^. Despite these remarkable advancements, current light-inducible systems tested and used in zebrafish suffer from low efficiency, leakiness in the dark, the need of prolonged exposure to UV light resulting in cytotoxicity, the requirement of advanced microscopy setups, the addition of exogenous chromophores, or the use of harmful chemicals such as caged biomolecules or tamoxifen^10,12–17^.

On the other hand, a highly efficient genetically encoded photoactivatable Cre recombinase (PA-Cre) has recently been reported in mammals based on Magnet dimerization systems^18–20^. The Magnet system was engineered from the blue light photoreceptor Vivid (VVD) which is derived from the filamentous fungus *Neurospora crassa.* and requires flavin adenine dinucleotide as its cofactor, which is abundant in eukaryotic cells^21,22^. This blue light-dependent dimerization system comprises two photoswitches named positive Magnet (pMag) and negative Magnet (nMag)^21^. Based on the reconstitution of split Cre fragments upon blue light-dependent dimerization of the Magnet system, Kawano et al. developed a genetically encoded Magnets-based PA-Cre system^18^. Consequently, they showed that their PA-Cre system could be applied to control DNA recombination with high spatiotemporal precision *in vivo*.

The Magnet system has also been shown to be efficient in inducing light-controlled genetic recombination *in vitro* and *in vivo* when used with Dre and Flp recombinases in mammals^23,24^. Furthermore, it has been recently reported that the Magnets-based photoactivatable split Gal4 system enables rapid and robust spatiotemporal gene expression control in *Drosophila* in vivo^25^. However, to date, there are no stable transgenic zebrafish models for optogenetic-based Cre/*loxP* recombination. Therefore, we aimed to develop a PA-Cre system to enable light-inducible lineage tracing *in vivo*, leveraging on the unique strengths of the zebrafish model.

In this study, we evaluated the functionality of the PA-Cre system in zebrafish. We demonstrate that the original PA-Cre system previously used in mammalian cells^18^, induces efficient Cre/*loxP* recombination in zebrafish embryos upon simple LED-mediated blue light illumination. We further developed and assessed the performance of the PA-Cre system under various zebrafish-specific promoters. We illustrate that our new system exhibits no phototoxicity or leakiness in the dark and enables efficient *loxP*-mediated DNA recombination *in vivo*. Finally, we generated a new transgenic zebrafish line in which the expression of the PA-Cre system is under the control of the zebrafish *ubiquitin* (*ubb*) promoter^26^. By utilizing this *Tg(ubb:PA-Cre)* line, we show that our system allows for efficient and robust Cre/*loxP* recombination in diverse tissues and cell types through noninvasive blue light illumination. Collectively, our newly developed versatile tool is anticipated to pave the way for light-controlled tracing of cell fate and migration *in vivo*, opening up new opportunities for zebrafish research.

## Results

### The PA-Cre system enables *in vivo loxP*-mediated DNA recombination in zebrafish embryos

To assess the efficacy of the PA-Cre system (Figure 1A) to induce *loxP*-mediated DNA recombination in zebrafish, we injected the original PA-Cre construct^18^ (Figure 1B) into zebrafish eggs. The PA-Cre construct, in which expression is driven by the cytomegalovirus (CMV) promoter^18^ (Figure 1B), was injected into *Tg(-3.5ubb:loxP-lacZ-loxP*-eGFP; *cry:GFP)* (referred to hereafter as *Tg(ubb:HULK)*^27^ embryos at one-cell stage and then subjected to global 488-nm blue light illumination with from 6 hours post-fertilization (hpf) to 18 hpf (Figure 1C) utilizing a simple low-cost LED light. It has been shown that The *Tg(ubb:HULK)* zebrafish exhibits ubiquitous green fluorescence protein (GFP) expression only upon Cre-mediated recombination^27^. In injected embryos, 488-nm blue light illumination at 6 hpf resulted in Cre/*loxP* recombination and generated a mosaic GFP expression in various tissues throughout the embryo body, including the head, the heart, the trunk and the tail (Figure 1D-E). All embryos that were injected and exposed to blue light illumination showed mosaic GFP expression. Importantly, we observed no GFP expression in embryos that were injected and kept in the dark (Figure 1D), indicating that the Cre-mediated recombination is solely driven by blue light illumination and does not undergo stochastic magnets dimerization. Fluorescence imaging showed that zebrafish embryos exposed to blue light exhibited a 29-fold induction over embryos that were kept in the dark (Figure 1F). In addition, we observed no photo-induced cytotoxicity upon 488-nm blue light illumination, as we found comparable rates of abnormalities between light-exposed embryos (13.3%) and non-exposed embryos (15.2%). These results demonstrate that transient photoactivation of the PA-Cre system in zebrafish embryos allows robust light-induced Cre/*loxP* recombination in multiple tissues. Importantly, no background DNA recombination was detected in the absence of illumination and no photo-induced cytotoxicity was observed.

**Figure 1:**
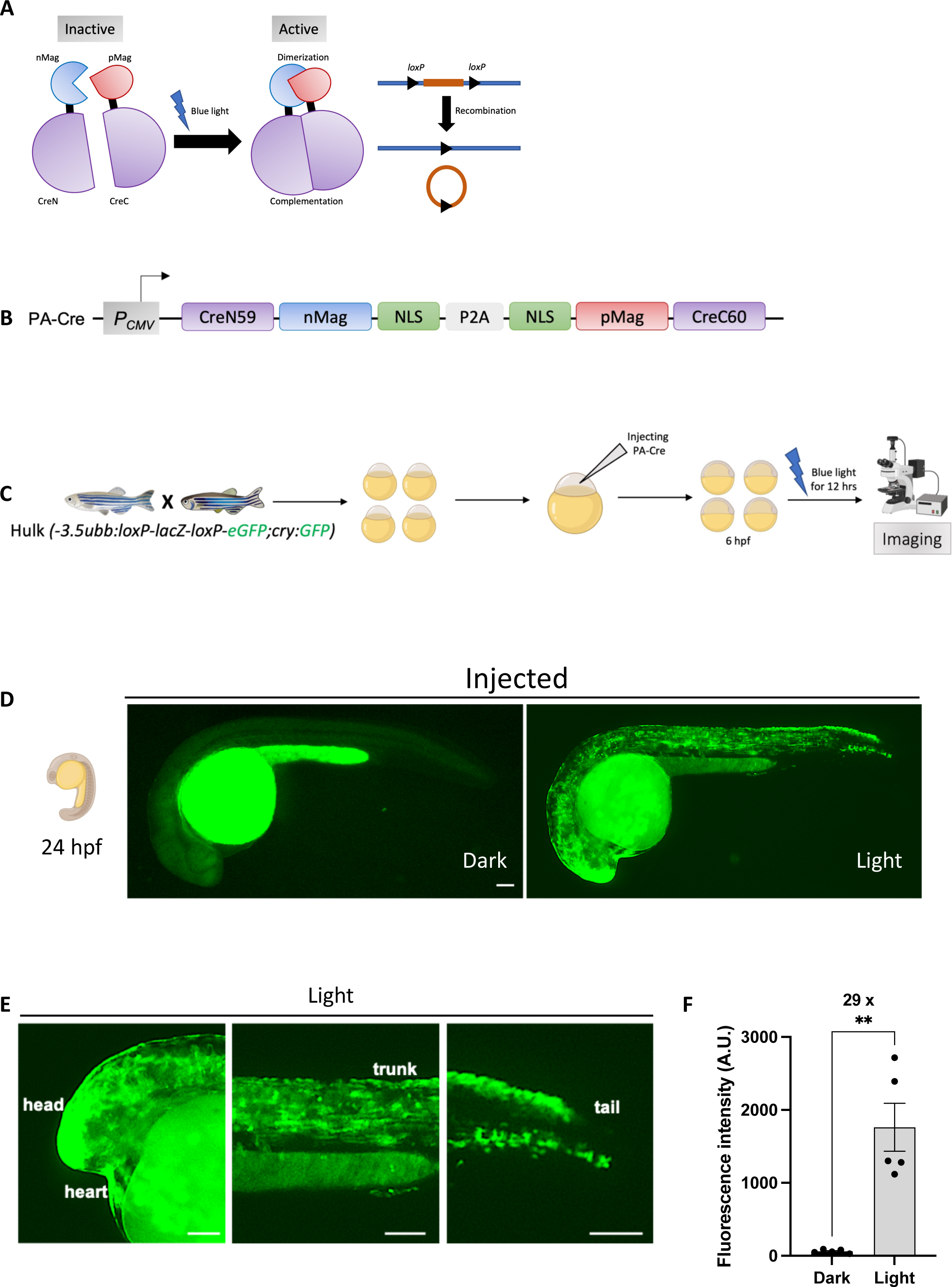
**PA-Cre mediated genetic recombination in zebrafish embryos.** (A) Schematic representation of the optogenetic PA-Cre system. CreN and CreC are respectively fused to nMagnet and pMagnet tandems. These fusions are monomers in the dark state (inactive PA-Cre). Upon blue light photoactivation, dimerization of nMag-pMag leads to complementation of CreN-CreC and thus induction of genetic recombination and activation of gene expression (active PA-Cre). nMag, negative Magnet; pMag, positive Magnet; CreN and CreC: two pairs of split Cre fragments. (B) Schematic representation of the PA-Cre genetic construct. The PA-Cre system is driven by the CMV promoter. P, promoter; nMag, negative Magnet; pMag, positive Magnet; NLS, nuclear localization signal; CreN59, split Cre fragment (residues 19–59); CreC60, split Cre fragment (residues 60–343); P2A, 2A self-cleaving peptides. (C) Schematic representation of the strategy used to induce genetic recombination in zebrafish embryos. *Tg(ubb:Hulk)* embryos were injected at the one-cell stage with the PA-Cre construct, photoactivated with blue light at 6 hpf for 12 hours, and then imaged at the desired developmental stage. (D) Representative fluorescent images at 24 hpf of injected *Tg(ubb:Hulk)* embryos kept in the dark (left panel) or exposed to blue light illumination at 6 hpf for 12 hours (right panel). Scale bar: 100 µm. (E) Closeups of image in (D) right panel showing GFP signal of a photoactivated 24 hpf embryo in the head, heart, trunk and tail. Scale bar: 100 µm. (F) Quantification of fluorescence intensity at 48 hpf of embryos injected with the PA-Cre construct and kept in the dark (n=5) or exposed to blue light illumination at 6 hpf for 12 hours (n=5). 29x represents the fold induction of the PA-Cre system. Mean ± SEM of the fluorescence intensity is shown. An unpaired t-test with Welch’s correction was used for this analysis. ∗∗p ≤ 0.01. A.U.;

### Generation of a stable functional zPA-Cre transgenic line in Zebrafish

Having established the functionality of the PA-Cre system is in zebrafish embryos, we tested its suitability for generating a stable transgenic line. To this end, we decided to use the zebrafish *ubb* promoter^26^ in place of the CMV promoter to control the expression of the PA-Cre system. The CMV promoter has been associated with potential silencing of promoter activity *in vivo*, making it less suitable for this type of applications. In contrast, the *ubb* promoter has been shown to have consistent activity throughout zebrafish development stages and does not exhibit silencing issues^26,28^. Therefore, we created the *Tg(-3.5ubb:PA-Cre; cmlc2-EGFP)* line (referred to hereafter as zPA-Cre) in which the expression of the PA-Cre system is driven by the *ubb* promoter. Additionally, the line features *cmlc2-EGFP* as a transgenesis marker in the heart (Figure 2A).

**Figure 2:**
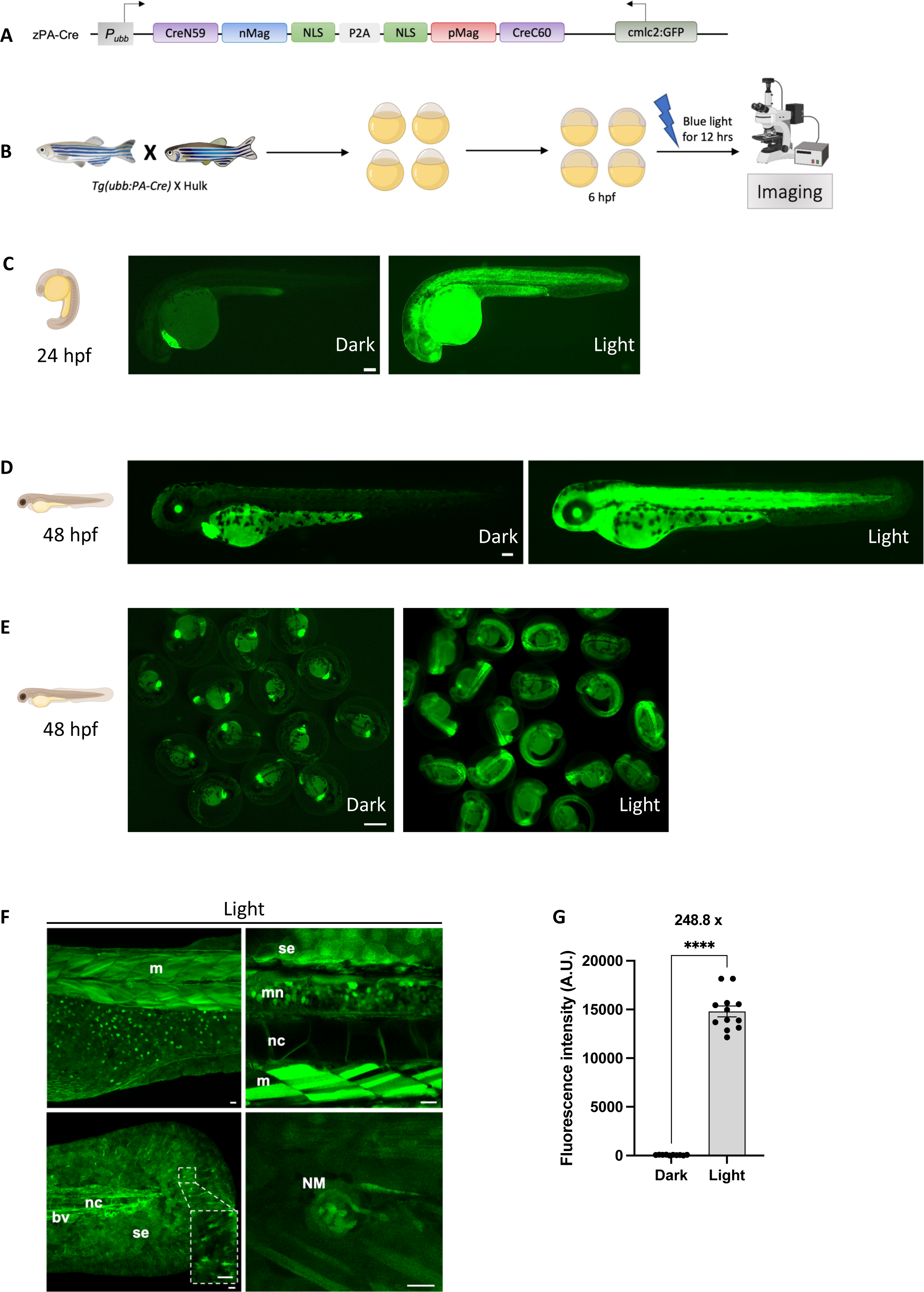
The stable transgenic line *Tg(ubb:PA-Cre)* enables light-induced genetic recombination in zebrafish embryos. (A) Schematic representation of the zebrafish PA-Cre genetic construct (zPA-Cre). The zPA-Cre system is driven by the ubb promoter. The green heart is used as a transgenesis marker (cmlc2:GFP). P, promoter; nMag, negative magnet; pMag, positive magnet; NLS, nuclear localization signal; CreN59, split Cre fragment (residues 19–59); CreC60, split Cre fragment (residues 60–343); P2A, 2A self-cleaving peptides; cmlc2, cardiac myosin light chain 2. (B) Schematic representation of the strategy used to induce genetic recombination in zebrafish embryos. The stable transgenic line *Tg(ubb:PA-Cre)* was crossed with *Tg(ubb:Hulk).* The double transgenic embryos *Tg(ubb:PA-Cre; ubb:Hulk)* were photoactivated with blue light at 6 hpf for 12 hours and then imaged at the desired developmental stage. (C) Representative fluorescent images at 24 hpf of *Tg(ubb:PA-Cre; ubb:Hulk)* embryos kept in the dark (left panel) or exposed to blue light illumination at 6 hpf for 12 hours (right panel). Scale bar, 100 µm. (D) Representative fluorescent images at 48 hpf of *Tg(ubb:PA-Cre; ubb:Hulk)* embryos kept in the dark (left panel) or exposed to blue light illumination (right panel) at 6 hpf for 12 hours. Scale bar, 100 µm. (E) Fluorescent images at 48 hpf of a group of *Tg(ubb:PA-Cre; ubb:Hulk)* embryos kept in the dark (left panel) or exposed to blue light illumination at 6 hpf for 12 hours (right panel). Scale bar, 100 µm. (F) Representative confocal fluorescent z-stack images of photoactivated *Tg(ubb:PA-Cre; ubb:Hulk)* embryo at 72 hpf. The higher magnification highlights GFP+ signal observed in various tissues and cells. m: skeletal muscle, se: skin epithelia, mn: motor neurons, nc: notochord, bv: blood vessels, NM: neuromast. Scale bar, 20 µm. (G) Quantification of fluorescence intensity at 48 hpf of *Tg(ubb:PA-Cre; ubb:Hulk)* embryos kept in the dark (n=10) or exposed to blue light illumination at 6 hpf for 12 hours (n=12). 248.8x represents the fold induction of the zPA-Cre system. Mean ± SEM of the fluorescence intensity is shown. Unpaired t-test with Welch’s correction was used for this analysis. ****p ≤ 0.0001. A.U.: Arbitrary Unit.

To test whether the newly developed line is capable of inducing DNA recombination upon 488-nm blue light illumination, we crossed it with the *Tg(ubb:HULK)* (Figure 2B). In double transgenic zebrafish embryos, blue light illumination from 6 hpf to 18 hpf resulted in a robust Cre/*loxP* recombination, leading to robust GFP expression in various tissues at 24 hpf (Figure 2C) and 48 hpf (Figure 2D-F). Confocal imaging revealed that embryos exposed to 488-nm blue light illumination showed GFP expression in several tissues and cell types, including skeletal muscle cells, the skin epithelia, motor neurons, the notochord, neuromasts and blood vessels (Figure 2F and supplemental video 1), while no GFP expression was observed in embryos kept in the dark (Figure 2C-E), confirming the fidelity of the zPA-Cre line and its inducibility exclusively upon blue light illumination. Furthermore, no photo-induced cytotoxicity was observed, as evidenced by the absence of Acridine Orange (AO)-positive cells throughout the body of the larvae exposed to blue light. In addition, there was no difference in the number of dead cells in the forebrain between light-exposed embryos and non-exposed embryos (supplemental Figure 1A-B). Furthermore, the abnormality rate of blue light exposed embryos (10.47%; n=219) was not considerably different from that of non-exposed embryos (10.28%; n=176). Moreover, fluorescence imaging demonstrated a 248.8-fold induction of GFP expression in zebrafish embryos exposed to blue light compared to those kept in the dark (Figure 2G). To assess how quickly GFP expression appears following exposure to light, we photoactivated *Tg(zPA-Cre; ubb:HULK)* zebrafish embryos at 1-cell stage. We observed that light-exposed embryos began expressing GFP at the 75% epiboly stage (8 hpf), and that GFP expression intensified over development (supplemental Figure 2A-B). Altogether, these findings demonstrate the robust and efficient induction of DNA recombination *in vivo* upon blue light illumination by the zPA-Cre line.

### zPA-Cre is effective in diverse cell types

To assess the versatility of the zPA-Cre system, we investigated its capacity to induce DNA recombination in various Cre-responsive transgenic zebrafish lines. We crossed the zPA-Cre line with distinct switch lines, in which specific cell types express DsRed under steady-state conditions and switch to GFP expression upon activation of Cre. These included the macrophage-specific *Tg(mpeg1:LoxP-DsRedx-LoxP-GFP-NTR)*^29^ (Figure 3A), or the erythrocyte-specific *Tg(α/βa2*-*globin:loxP-DsRedx-loxP-GFP)*^30^ line (Figure 3B) or the *lymphocyte-specific Tg(lck:loxP-DsRedx-loxP-GFP)*^30,31^ line (Figure 3C). In double transgenic zebrafish embryos, exposure to 488-nm blue light illumination from 6 hpf to 18 hpf resulted in Cre/*loxP*, leading to robust GFP expression at 48 hpf in macrophages (Figure 3A) and in erythrocytes (Figure 3B), and at 120 hpf in thymocytes (Figure 3C). To evaluate the efficiency of the zPA-Cre system, we quantified the proportion of switched GFP-positive macrophages, erythrocytes and thymocytes in the *Tg(mpeg1:LoxP-DsRedx-LoxP-GFP-NTR)* line, the *Tg(α/βa2*-*globin:loxP-DsRedx-loxP-GFP)* line and the *Tg(lck:loxP-DsRedx-loxP-GFP)* line, respectively. We found that ∼95% of macrophages (Figure 3D), *∼*90% of erythrocytes (Figure 3E) and ∼82% of thymocytes in the thymus (Figure 3F) exhibited GFP expression, indicating the high efficiency of our labelling system.

**Figure 3:**
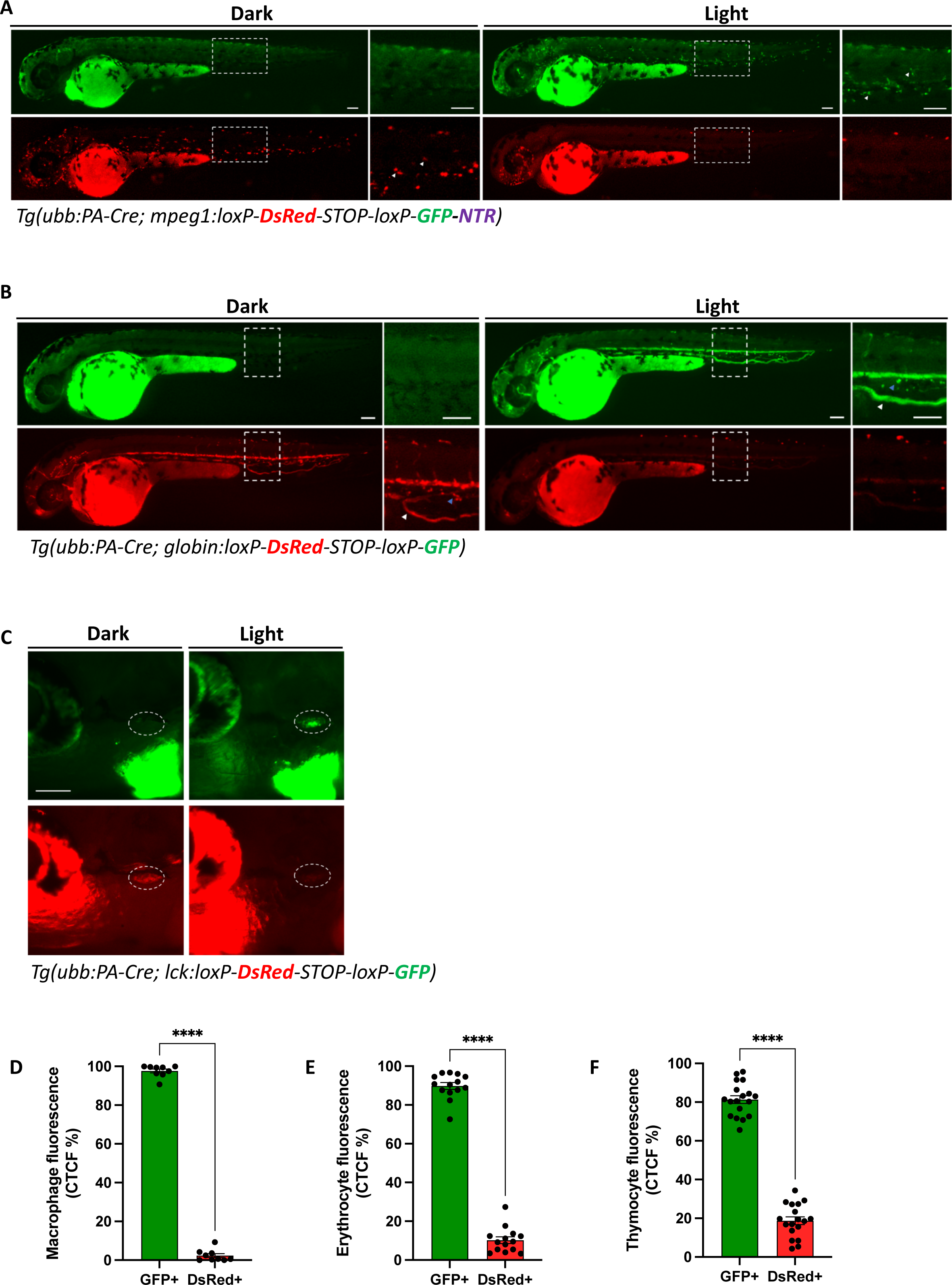
***Tg(ubb:PA-Cre)* enables light-induced genetic recombination with different Cre reporter zebrafish transgenic lines.** (A) Representative fluorescent images at 48 hpf of *Tg(ubb:PA-Cre; mpeg1:LoxP-DsRedx-LoxP-GFP-NTR)* embryos kept in the dark or exposed to blue light illumination at 6 hpf for 12 hours. White dashes indicate the region with higher magnification. White arrowheads highlight macrophages at higher magnification in the trunk. Scale bar, 100 µm. (B) Representative fluorescent images at 48 hpf of *Tg(ubb:PA-Cre; α/βa2-globin:loxP-DsRedx-loxP-GFP)* embryos kept in the dark or exposed to blue light illumination at 6 hpf for 12 hours. White dashes indicate the region with closeups. White arrowheads highlight erythrocytes in circulation, and blue arrowheads highlight erythrocytes in the CHT. Scale bar, 100 µm. (C) Representative fluorescent images at 120 hpf of *Tg(ubb:PA-Cre; lck:loxP-DsRedx-loxP-GFP)* embryos kept in the dark or exposed to blue light illumination at 6 hpf for 12 hours. White dashes mark the thymus. Scale bar, 100 µm. (D) Quantification of macrophage fluorescence intensity was measured at 48 hpf in photoactivated *Tg(ubb:PA-Cre; mpeg1:Switch)* embryos (n=9). Mean ± SEM of the DsRed+ and GFP+ corrected total cell fluorescence (CTCF) percentage is shown. Unpaired t-test with Welch’s correction was used for this analysis. ****p ≤ 0.0001. (E) Quantification of erythrocyte fluorescence intensity was measured at 48 hpf in photoactivated *Tg(ubb:PA-Cre; globin:Switch)* embryos (n=14). Mean ± SEM of the DsRed+ and GFP+ corrected total cell fluorescence (CTCF) percentage is shown. Unpaired t-test with Welch’s correction was used for this analysis. ****p ≤ 0.0001. (F) Quantification of thymocyte fluorescence intensity was measured at 120 hpf in photoactivated *Tg(ubb:PA-Cre; lck:Switch)* embryos (n=18). Mean ± SEM of the DsRed+ and GFP+ corrected total cell fluorescence (CTCF) percentage is shown. Unpaired t-test with Welch’s correction was used for this analysis. ****p ≤ 0.0001.

In addition, we tested the functionality of the PA-Cre system under the *kdrl* promoter^32,33^, which exhibits expression during gastrulation, in the lateral plate mesoderm, and in endothelial cells^33–35^. To examine the potential of the *kdrl*-PA-Cre system for erythrocyte labeling, we injected a *kdrl*-PA-Cre construct into *Tg(α/βa2*-*globin:loxP-DsRedx-loxP-GFP)* embryos at one-cell stage, as erythrocytes are known to be derived from the lateral plate mesoderm^36–39^. Following blue light illumination from 6 hpf to 18 hpf in injected embryos, GFP expression was observed in erythrocytes at 24 hpf, indicating successful Cre/loxP recombination. Conversely, non-exposed embryos did not exhibit GFP expression in erythrocytes (supplemental Figure 3A-B). These results indicate that the zPA-cre system enables DNA recombination in different cell types, and that the PA-Cre system expression can be driven by distinct promoters. Collectively, our data establish the zPA-Cre system as a versatile tool for DNA recombination in zebrafish.

### zPA-Cre enables light-inducible Cre/*loxP* recombination at different developmental time points

We then evaluated the efficiency of our system in comparison to widely used conventional non-photoactivatable Cre lines. To this end, we utilized the *Tg(ubb:creERT2)* line, in which Cre is ubiquitously expressed but its activity is induced only by incubation with tamoxifen, that allows Cre to enter to the nucleus^26^. We found that *Tg(zPA-Cre; ubb:HULK)* zebrafish embryos exposed to blue light overnight at 6 hpf exhibited a 23.2-fold induction compared to *Tg(ubb:creERT2; ubb:HULK)* embryos treated with 4-OH-tamoxifen (4-OHT) (Figure 4A,C), indicating that our system exhibits superior induction efficiency compared to other conventional inducible tools.

**Figure 4:**
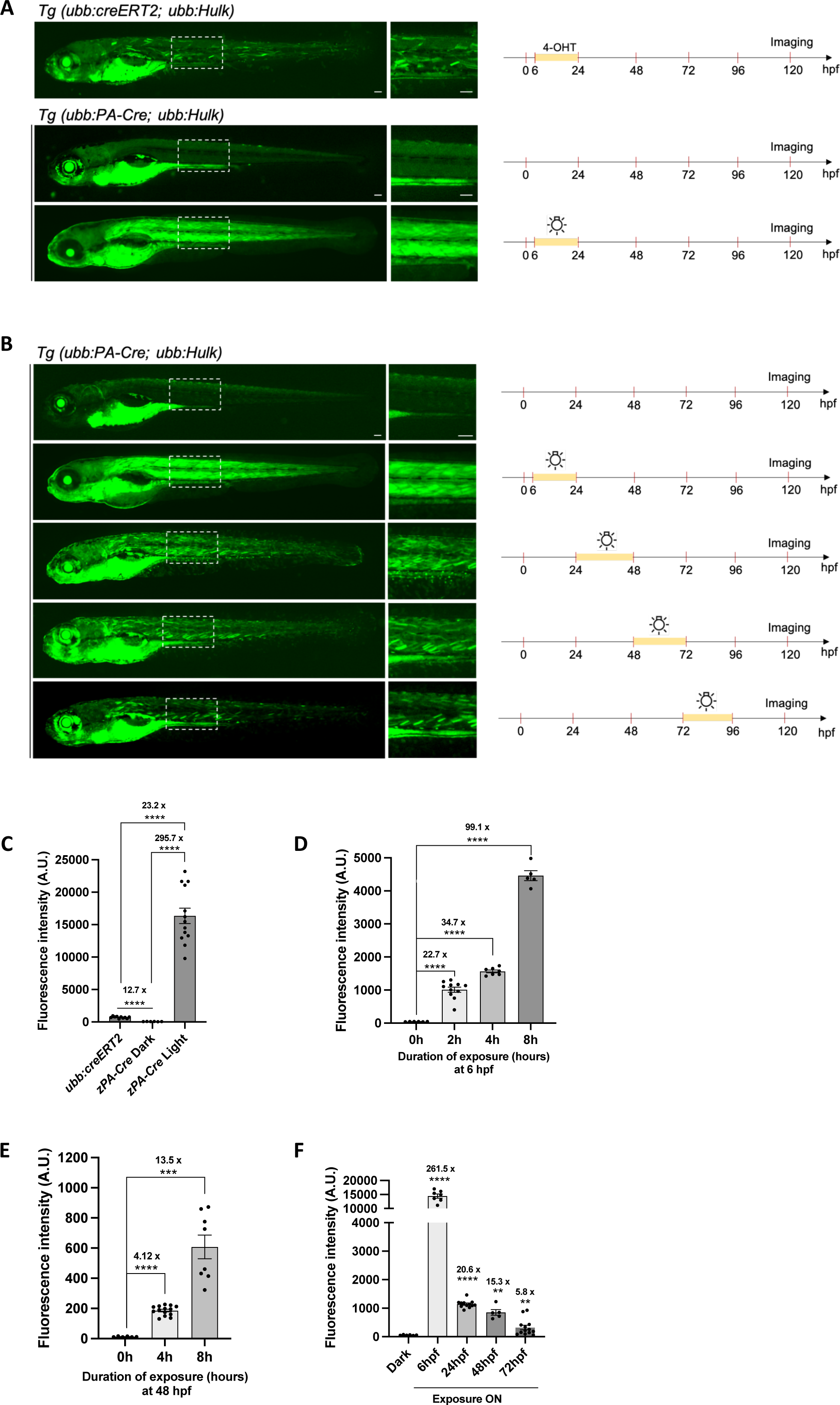
***Tg(ubb:PA-Cre)* enables light-induced Cre-LoxP recombination at different developmental stages** (A) Representative fluorescent images at 120 hpf of 4-OHT-induced *Tg(ubb:creERT2; ubb:Hulk)* embryo or *Tg (ubb:PA-Cre; ubb:Hulk)* embryo kept in the dark or exposed to blue light illumination at 6 hpf overnight (left panel). White dashes indicate the region showing closeups. Scale bar, 100 µm. Schematic representation of the strategy used to induce genetic recombination in zebrafish *Tg(ubb:creERT2; ubb:Hulk)* or *Tg(ubb:PA-Cre; ubb:Hulk)* embryos (right panel). (B) Representative fluorescent images at 120 hpf of *Tg(ubb:PA-Cre; ubb:Hulk)* embryos kept in the dark or exposed to blue light illumination overnight at 6 hpf, 24 hpf, 48 hpf, and 72 hpf (left panel). White dashes indicate the region with closeups. Scale bar, 100 µm. Schematic representation of the strategy used to induce genetic recombination in zebrafish *Tg(ubb:PA-Cre; ubb:Hulk)* embryos at different developmental stages (right panel). (C) Quantification of fluorescence intensity at 120 hpf of 4-OHT-induced *Tg(ubb:creERT2; ubb:Hulk)* (n=10), *Tg(ubb:PA-Cre; ubb:Hulk)* embryo kept in the dark (n=6), or exposed to blue light illumination at 6 hpf overnight (n=13). The numbers 23.2x and 295.7x represent the fold induction comparison between the light-exposed group and the 4-OHT-induced group, and between the light-exposed group and the non-exposed ones, respectively. Mean ± SEM of the fluorescence intensity is shown. Unpaired t-test with Welch’s correction was used for this analysis. ****p ≤ 0.0001. A.U., Arbitrary Unit. (D) Quantification of fluorescence intensity at 120 hpf of *Tg(ubb:PA-Cre; ubb:Hulk)* embryos exposed to blue light illumination at 6 hpf for 0 hours (n=6), 2 hours (n=11), 4 hours (n=7), and 8 hours (n=5). The numbers 22.7x, 34.7x, and 99.1x represent the fold induction comparison between the 2-hour exposed group and the non-exposed group, the 4-hour exposed group and the non-exposed group, and the 8-hour exposed group and the non-exposed group, respectively. Mean ± SEM of the fluorescence intensity is shown. Unpaired t-test with Welch’s correction was used for this analysis. ****p ≤ 0.0001. A.U., Arbitrary Unit. (E) Quantification of fluorescence intensity at 120 hpf of *Tg(ubb:PA-Cre; ubb:Hulk)* embryos exposed to blue light illumination at 48 hpf for 0 hours (n=6), 4 hours (n=14), and 8 hours (n=8). The numbers 4.12x and 13.5x represent the fold induction comparison between the 4-hour exposed group and the non-exposed group, and the 8-hour exposed group and the non-exposed group, respectively. Mean ± SEM of the fluorescence intensity is shown. Unpaired t-test with Welch’s correction was used for this analysis. ***p ≤ 0.001; ****p ≤ 0.0001. A.U., Arbitrary Unit. (F) Quantification of fluorescence intensity at 120 hpf of *Tg(ubb:PA-Cre; ubb:Hulk*) embryos kept in the dark (n=6) or exposed to blue light illumination overnight at 6 hpf (n=7), at 24 hpf (n=12), at 48 hpf (n=5), and at 72 hpf (n=13). The numbers 261.5x, 20.6x, 15.3x, and 5.8x represent the fold induction comparison between the different groups and the dark one, respectively. Mean ± SEM of the fluorescence intensity is shown. Unpaired t-test with Welch’s correction was used for this analysis. **p ≤ 0.01; ****p ≤ 0.0001. A.U., Arbitrary Unit.

The use of light-inducible tools in zebrafish beyond the embryonic stages can be challenging due to limited light penetration in larval tissues. Therefore, we assessed the efficiency of DNA recombination using the zPA-Cre system at different developmental time points. To this end, we exposed double transgenic *Tg(zPA-Cre; ubb:HULK)* embryos to blue light overnight at 24 hpf, 48 hpf, and 72 hpf, and imaged them at 120 hpf (Figure 4B,F). We observed GFP expression in all embryos or larvae exposed to blue light, although at lower levels compared to embryos exposed to blue light at 6 hpf. Of note, control siblings that were kept in the dark did not show GFP expression (Figure 4B, F). Next, we investigated the relationship between the zPA-Cre activity and the duration of blue light illumination. We exposed double transgenic *Tg(zPA-Cre; ubb:HULK)* embryos to blue light for various time periods at 6 hpf and 48 hpf. We observed that longer exposure times positively correlated with increased GFP expression in embryos. Notably, illuminating 6 hpf embryos with blue light for only 2 hours resulted in robust GFP expression at 120 hpf (Figure 4D,E and supplemental Figure 4A). These results indicate that zPA-Cre enables efficient DNA recombination even with short periods of blue light exposure, and that the zPA-Cre activity increases in a light dose-dependent manner.

Furthermore, to assess the applicability of our zPA-Cre system in inducing DNA recombination beyond embryonic stages, we photoactivated double transgenic *Tg(zPA-Cre; ubb:HULK)* 6 dpf larvae. We found that the zPA-Cre line can be utilized to induce expression at larval stages, thus extending the use of this system beyond early embryonic stages (supplemental Figure 4B-D). These findings demonstrate that while the highest efficiency is achieved during early developmental stages, our system remains effective at later developmental stages.

### zPA-Cre enables optogenetic cell ablation in zebrafish embryos

One of the challenges in characterizing the role of specific population subsets is the lack of tools for cell type-specific ablation^40^. Despite the widespread use of chemically inducible Cre strains to achieve this goal, the temporal and spatial resolution is limited. To demonstrate the ability to perform light-induced cell ablation, zPA-Cre fish were crossed with the macrophage-specific *Tg(mpeg1:LoxP-DsRedx-LoxP-GFP-NTR)*, allowing selective ablation of macrophages based on their origin. In this line, the expression of bacterial nitroreductase (NTR) is under the control of the *mpeg1* promoter^29^. Thus, NTR will be expressed exclusively in the photoactivated macrophages; consequently, these macrophages could be ablated by metronidazole (MTZ) treatment^41^ (Figure 5A). First, we photoactivated double transgenic *Tg(zPA-Cre; mpeg1:LoxP-DsRedx-LoxP-GFP-NTR)* between 14 hpf and 24 hpf, a time when primitive macrophages start to emerge in the developing embryo. To evaluate the efficiency of our labelling strategy, we quantified the proportion of switched GFP-positive macrophages and found that ∼65% of macrophages exhibited GFP expression (Figure 5C), indicating the high efficiency of our labelling strategy.

**Figure 5:**
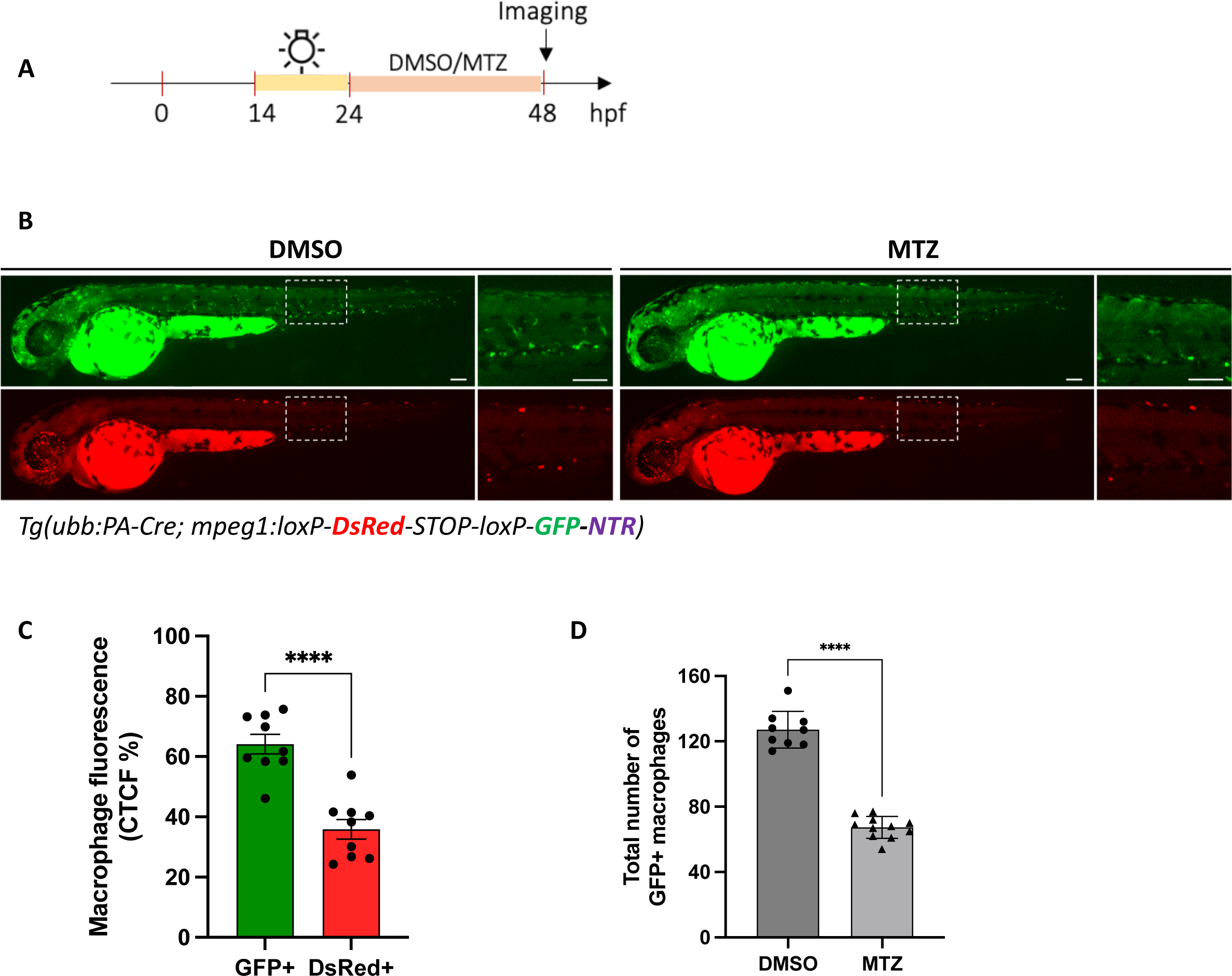
***Tg(ubb:PA-Cre)* enables optogenetic cell ablation in zebrafish embryos** (A) Schematic representation strategy used to selectively ablate macrophages in photoactivated embryos. (B) Representative fluorescent images at 48 hpf of *Tg(ubb:PA-Cre; mpeg1:LoxP-DsRedx-LoxP-GFP-NTR)* embryos treated with either DMSO as control, or metronidazole (MTZ) to ablate macrophages. White dashes indicate the region showing closeups. Scale bar, 100 µm. (C) Quantification of macrophage fluorescence intensity was measured at 48 hpf in photoactivated *Tg(ubb:PA-Cre; mpeg1:LoxP-DsRedx-LoxP-GFP-NTR)* embryos treated with DMSO (n=9). Mean ± SEM of the DsRed+ and GFP+ corrected total cell fluorescence (CTCF) percentage is shown. Unpaired t-test with Welch’s correction was used for this analysis. ****p ≤ 0.0001. (D) Quantification of the total number of GFP+ macrophages at 48 hpf in photoactivated *Tg(ubb:PA-Cre; mpeg1:LoxP-DsRedx-LoxP-GFP-NTR)* embryos treated with either DMSO (n=9) or MTZ (n=11). Unpaired t-test with Welch’s correction was used for this analysis. ****p ≤ 0.0001.

To assess the efficiency of the zPA-Cre-mediated cell ablation with the NTR-MTZ system, we quantified the total number of GFP+ macrophages in photoactivated embryos after treatment with either DMSO or MTZ for 24 hours from 24 to 48 hpf. We observed a significant reduction of macrophages with MTZ treatment compared to the DMSO-treated controls, suggesting that zPA-Cre also enables light-induced cell type-specific ablation (Figure 5B and D). In summary, our zPA-Cre transgenic line can be utilized for efficient spatiotemporal cell ablation in various tissues, providing a valuable approach for dissecting cell type-specific functions.

### zPA-Cre enables light-inducible Cre/*loxP* recombination at a specific region of interest and at single-cell resolution

To test the capability of the zPA-Cre system to drive DNA recombination in specific tissue regions, we used the *Tg(zPA-Cre; ubb:HULK)* embryos and performed photoactivation in targeted regions of interest. Photoactivation with blue light using a 488 nm laser at 100% laser power (2 mW/mm2) in a specific region (a 25 um diameter circle) for 5 minutes within a single skeletal muscle at 48 hpf resulted in GFP expression in the activated region (Figure 6A).

**Figure 6:**
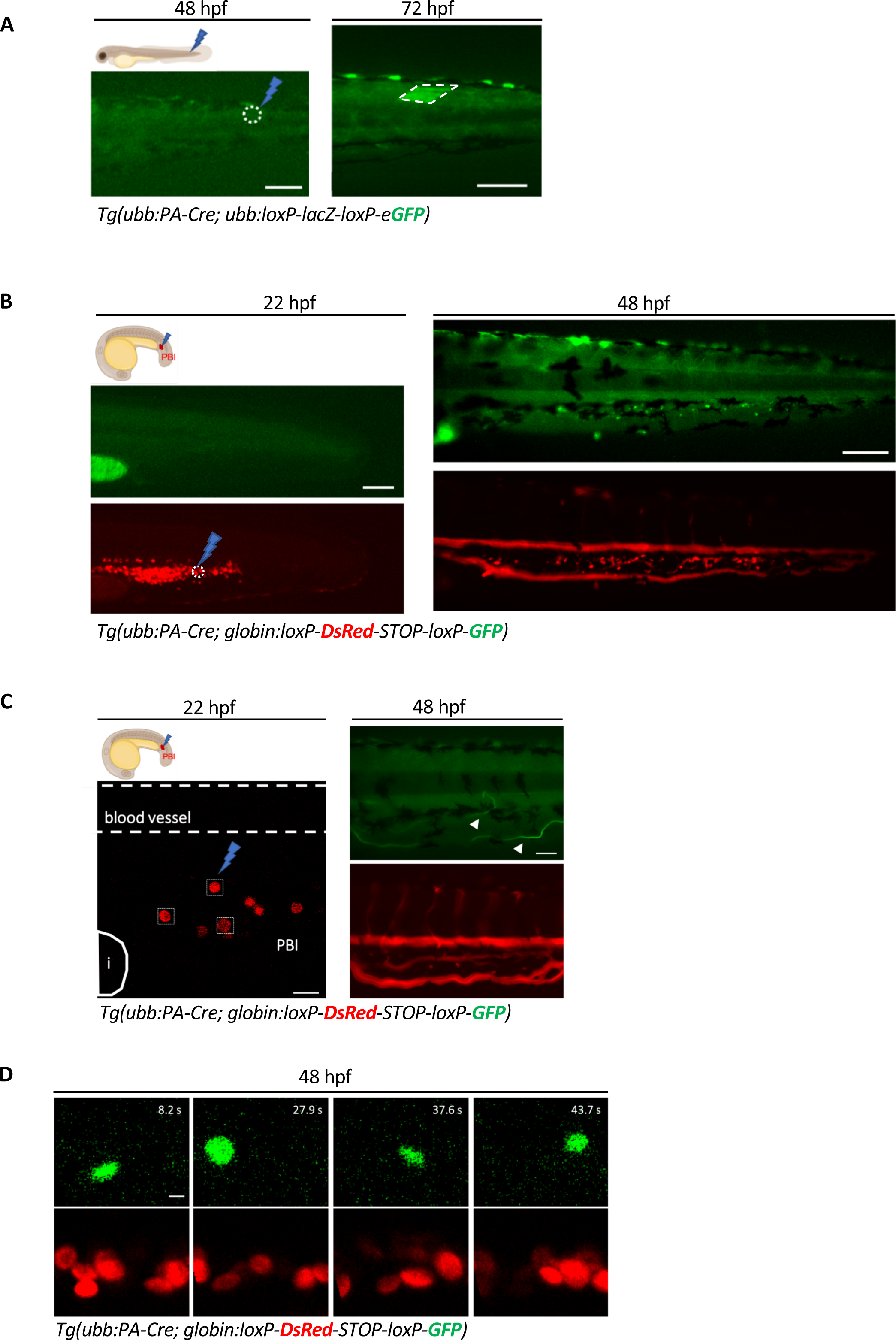
***Tg(ubb:PA-Cre)* enables light-induced spatial and temporal genetic recombination.** (A) Schematic representation strategy used to induce local photoactivation within a single skeletal muscle cell using a blue light laser. A 25 µm diameter circle within a single skeletal muscle was targeted by a 488 nm laser for 5 minutes (n=6). Scale bar, 100 µm. Representative fluorescent images at 72 hpf of *Tg(ubb:PA-Cre; ubb:Hulk)* showing GFP+ skeletal muscle. Scale bar, 100 µm. (B) Schematic representation strategy used to induce local photoactivation with blue light laser at 22 hpf. Few cells in the PBI (a 25 μm diameter circle) were targeted by a 488 nm laser for 5 minutes (n=6; left panel). Scale bar, 100 µm. Representative fluorescent image at 48 hpf of *Tg(ubb:PA-Cre; α/βa2-globin:loxP-DsRedx-loxP-GFP)* embryo showing GFP+ erythrocytes in the CHT (right panel). Scale bar, 100 µm. (C) Schematic representation strategy used to induce two-photon photoactivation at 22 hpf. An area of 8x8 µm in the PBI that covers single erythroid progenitors was targeted by a 920 nm two-photon laser for 45 seconds (n=9; left panel). Scale bar, 20 µm. Representative fluorescent images at 48 hpf of *Tg(ubb:PA-Cre; α/βa2-globin:loxP-DsRedx-loxP-GFP)* showing GFP+ erythrocytes (arrowheads) in the circulation. Scale bar, 50 µm. Higher magnification images are shown in D. (D) Higher magnification images of still fluorescent images of time-lapse imaging in *Tg(ubb:PA-Cre; α/βa2-globin:loxP-DsRedx-loxP-GFP)* at 48 hpf showing GFP+ erythrocytes in the circulation. Scale bar, 5 µm.

Next, we utilized the *Tg(zPA-Cre)* line in combination with the *Tg(α/βa2*-*globin:loxP-DsRedx-loxP-GFP)* line to test if *loxp*-mediated DNA recombination could be induced in a small number of few erythrocytes present in the posterior blood island (PBI), a transient hematopoietic tissue in zebrafish^37^. Photoactivation with blue light using a 488 nm laser at 100% laser power (2 mW/mm2) in few erythrocytes in the PBI (with a 25 um diameter circle) for 5 minutes at 22 hpf resulted in the presence of GFP^+^ erythrocytes in the caudal hematopoietic tissue (CHT), the fetal liver counterpart in zebrafish^9^ (Figure 6B), as well as in the circulation (supplemental video 2) at 48 hpf. Finally, we examined if the zPA-Cre allows for optogenetic manipulation with single-cell resolution. To this end, we applied two-photon 920-nm illumination on the double *Tg(zPA-Cre; α/βa2*-*globin:loxP-DsRedx-loxP-GFP)* transgenic line to target single erythrocytes in the PBI region at 22 hpf, before the onset of blood circulation in zebrafish^37^ (Figure 6C). We applied 45 seconds of 920-nm illumination by scanning the infrared laser beam over an area of 8 um x 8 um to cover single erythroid progenitors. After photo-activation, embryos were kept in the dark for 24 hours, and live imaging was conducted at this stage to trace the migration of the photoactivated GFP+ cells and their progeny. We found that two-photon photoactivation of single erythrocytes resulted in the presence of GFP+ erythrocytes in the circulation, demonstrating that our zPA-Cre system can be used to trace the progenies derived from a single cell (Figure 6C-D and supplemental Video 3A-B).

In conclusion, these experiments demonstrate that the zPA-Cre system enables efficient and precise spatial and temporal control of *loxP*-mediated DNA recombination in zebrafish embryos with high resolution.

## Discussion

Optogenetic tools play a crucial role in studying development and tissue regeneration in various organisms, offering precise spatial and temporal resolution. Numerous light-inducible tools for Cre/*loxP* recombination have been developed utilizing UV light, blue light, or red light. Consequently, several attempts have been made to develop optically controlled Cre systems in zebrafish, as reported in previous studies^12–16^. Despite significant advances, current light-inducible systems tested in zebrafish have been performed in transient, and suffer from low efficiency, leakiness, require long light exposure resulting in cytotoxicity, require exogenous chromophores or need harmful chemicals as caged biomolecules or tamoxifen ^10,12–17^.

Here, we demonstrated the successful development of a novel stable optogenetic zebrafish line in which the PA-Cre system is controlled by the zebrafish *ubiquitin* promoter^26^. By utilizing the *Tg(ubb:PA-Cre)* line, we provide evidence that our system allows for efficient and robust Cre/*loxP* recombination in various tissues and cell types through noninvasive blue light illumination.

As a general rule, it is crucial for a light-inducible Cre recombinase to maintain low levels of background recombination^10^. In this regard, we show that the *Tg(ubb:PA-Cre)* line exhibits no background recombination in the absence of light stimulation, while displaying a highly inducible efficiency of DNA recombination upon exposure to blue light. Furthermore, we showcase the versatility of our system by demonstrating its ability to induce recombination at different developmental stages and achieve spatial control within specific regions upon blue light illumination. Importantly, after two-photon light illumination, the *Tg(ubb:PA-Cre)* line allows for optogenetic manipulation with single-cell resolution. These distinctive advantages offered by the *Tg(ubb:PA-Cre)* line pave the way for *in vivo* zebrafish studies during development with high spatiotemporal resolution.

As shown in this study, the newly generated *Tg(ubb:PA-Cre)* line enables precise spatial and temporal control of Cre/*loxP* recombination through blue light illumination. This versatile DNA recombination system holds potential for applications such as fate mapping or the induction of carcinogenesis in specific cell types upon illumination. Moreover, the utilization of the PA-Cre system in zebrafish reveals a striking simplicity and efficiency, thus expanding the repertoire of available genetic manipulation methods.

Our newly developed versatile tool is expected to pave the way for light-controlled DNA recombination *in vivo* in cells or tissues that may not have a specific promoter, thereby facilitating studies on lineage tracing, development, and disease. Furthermore, as we generated a stable transgenic line in which the PA-Cre system is driven by the *ubb* promoter, it holds a particular promise for investigating the utility of our tool in adult zebrafish. Exploring the performance and efficiency of the PA-Cre system under different promoters in zebrafish is also an enticing avenue to achieve more precise cell-specific DNA recombination. Lastly, with the development of the PA-Dre^23^ and PA-Flp^24^ systems, it will be become feasible to build additional optogenetic tools in zebrafish and perform intersectional genetics involving more than one recombinase.

Taken together, our findings indicate that the *Tg(ubb:PA-Cre)* line is an efficient optogenetic tool for gene manipulation with greatly expandable applications in developmental biology research.

### Limitations of the study

In this study, we achieved Cre/loxP recombination by either global blue light exposure or through laser-mediated illumination within a defined region or at the single-cell level. However, our zPA-Cre system is driven by the zebrafish *ubb* promoter which is active in various tissues. Further work is needed to assess whether the magnet split-Cre system can be expressed by tissue- or cell type-specific promoters to enable Cre/loxP recombination upon global blue light exposure in a tissue-specific manner. Additionally, it is necessary to test whether the two elements of the magnet split-Cre system can be expressed by two distinct promoters to allow for higher lineage tracing specificity. While we have demonstrated the utility of our system during early zebrafish developmental stages, further work is needed to assess whether this zPA-Cre system is also applicable to inducing DNA recombination in juvenile and adult zebrafish.

## STAR★Methods

### Key resources table

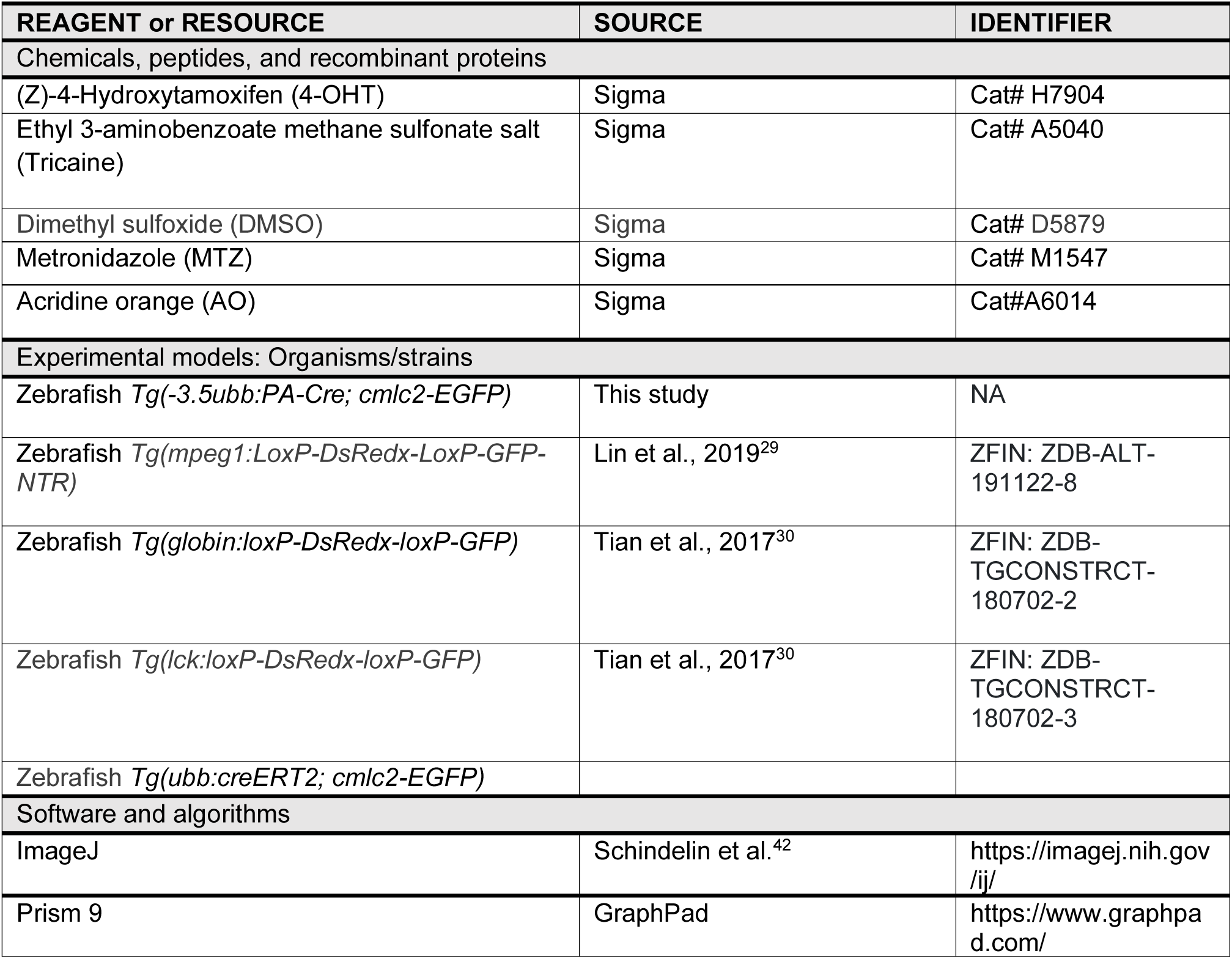

## Resource availability

### Lead contact

Further information and requests for scripts, resources, and reagents should be directed to and will be fulfilled by lead contact, Pedro P. Hernández.

### Data and code availability

This paper does not report original code. Any additional information required to reanalyze the data reported in this paper is available from the corresponding authors upon request.

### Materials and Data availability

The new reagents and data sets generated during the current study are available from the corresponding authors upon request.

## Materials and Methods

### Zebrafish

Embryonic, larval and adult zebrafish (*Danio rerio*) were maintained at 28°C on a 14-hour light/10-hour dark cycle. The collected embryos were raised in fish water containing 0.01% methylene blue to prevent fungal growth. All fish are housed in the fish facility of our laboratory, which was built according to the local animal welfare standards. All animal procedures were performed in accordance with French and European Union animal welfare guidelines and were approved by the ethic committee of the Curie Institute (Paris, France).

### Transgenic lines

The following lines of the AB stain were used*: Tg(ubb:HULK)*^27^*, Tg(mpeg1:LoxP-DsRedx-LoxP-GFP-NTR)*^29^*,Tg(globin:loxP-DsRedx-loxP-GFP)*^30^, *Tg(ubb:CreERT2; cmlc2-GFP)*^26^ and *Tg(lck:loxP-DsRedx-loxP-GFP)*^30^. All fish used in this study were homozygous and were crossed to produce embryos and larvae by natural mating. Embryo and larval zebrafish were studied before the onset of sexual differentiation and their sex can therefore not be determined.

### Plasmids

The original PA-Cre construct driven by the CMV promoter was purchased from Addgene (Addgene no. 122960). The PA-Cre coding sequence was synthesized by GenScript and listed in the supplementary note. The (*-3.5ubb:PA-Cre; cmlc2-EGFP)* constructs were generated by inserting 3.5 kb ubi promoter fragment, together with the coding sequence of PA-Cre, via Gibson assembly into pTol2 vector that carries cmlc2-EGFP as a transgenesis marker. To generate the *kdrl:PA-Cre* constructs, the *kdrl* promoter and the coding sequence of PA-Cre were cloned via Gibson assembly into the pTol2 vector.

### Zebrafish embryo microinjection

The *CMV:PA-Cre* and *kdrl:PA-Cre* DNA constructs (25 ng/μl) were injected into zebrafish embryos at the one cell stage using pressure-controlled microinjector and a micromanipulator. After injection, the embryos were cultured in E3 embryo medium with methylene blue in the dark at 28.5°C.

### Acridine Orange (AO) Staining

Zebrafish embryos were incubated in 5ug/ml AO solution for 1 hour and washed for several times until the background is clear.

### Generation of transgenic lines

The *-3.5ubb:PA-Cre;cmlc2-EGFP* constructs (25 ng/μl) were then co-injected with mRNA transposase (50 ng/μl) into 1-cell stage WT embryos^43^ and the resulting embryos were grown to adulthood for stable line screening. F0 founders were screened for specific *cmlc2*:GFP expression, raised to adulthood, and screened for germline transmission. Single-insertion transgenic strains were established and verified through screening for a 50% germline transmission rate in outcrosses in subsequent generations^44^.

### Measurements of Laser power

Single photon lasers: Laser powers were measured on the front focal plane of the objective using a compact power and energy meter console (ThorLabs, #PM100D) fitted with a low power photodiode sensor (ThorLabs, #S170C). Infrared multiphoton laser: Laser power on the focal plane was determined by measuring the power at the back focal plane of the objective with a thermopile sensor (PM10V1, Coherent, Santa Clara, USA) and multiplying by a factor of 0.75 (transmission of the 40x oil objective at 920 nm according to Zeiss transmission curves https://www.micro-shop.zeiss.com/) for the two-photon Ti:Sapphire laser used in this study.

### Photoactivation with global blue light

Global light induction was performed using a MARS AQUA-YWT-165W LED aquarium hood. Actual power of light exposure received by embryos, with lids of plates removed, was measured as ∼1.8mW/cm2 at 488 nm. 6 hpf embryos were illuminated with constant blue light for 12 h. To ensure all cells were exposed, embryos were periodically swirled during irradiation. In experiments presented in Figure 4, embryos and larvae at each indicated developmental stage were illuminated with constant blue light overnight and kept again in the dark till the desired developmental stage. Dark controls were placed in a lightproof box within the same 28.5°C incubator as the light-treated samples.

### Macrophage ablation and drug treatments

The *Tg(ubb:PA-Cre;mpeg1:loxP-DsRedx-loxP-GFP-NTR)* embryos were photoactivated by global blue light illumination and were immersed in system water containing 10 mM metronidazole (M1547, Sigma) for 24 hours, which caused an acute depletion of GFP-NTR^+^ cells. Photoactivated controls were incubated in the equivalent amount of DMSO solution during the same period. Light exposure was avoided by using foil to cover the plates as MTZ is light sensitive. Zebrafish were further analyzed by fluorescence imaging at 48 hpf.

### Regional activation

The blue-light activation was performed on a Zeiss LSM780 inverted confocal microscope. The desired region of the embryo was focused under a 25X lens (Zeiss Plan-Apochromat 25×, NA = 0.8 WD = 0.55), with the background fluorescence signal excited by a 405 nm laser. The desired region was selected under the bleach mode in ZEN software. A 2 mW 488 nm laser was used for activation the region for 5 minutes at 100% laser power. After activation, the embryos were incubated in dark at 28.5°C and then examined at indicated developmental stages.

### Two photon photoactivation for single cell targeting

Two-photon-based photoactivation was performed on a Zeiss LSM880 NLO inverted confocal microscope equipped with Ti:Sapphire laser (Mai Tai DeepSee, Spectra Physics) at 920 nm. The desired region of the embryo was focused under a 40X lens (Zeiss Plan-Apochromat 25×, NA = 1.3 WD = 0.21), with the red fluorescence signal excited by a 561 nm laser. The desired region was selected under the bleach mode in ZEN software. A 120 mW 920 nm laser was used for activation of single cells for 45 seconds. After activation, the embryos were incubated in dark at 28.5°C and then examined at indicated developmental stages.

### Fluorescence Microscopy

Zebrafish embryos, larvae and adults were anesthetized with 0.01% tricaine (A5040, Sigma) and mounted in 2.5% methylcellulose in 35-mm imaging dishes (MatTek) as described previously^45^. Fluorescent imaging was performed with either Zeiss LSM780 inverted confocal microscope or Zeiss Axio Zoom V.16 upright microscope with an AxioCam HRm Zeiss camera and Zeiss Zen 3.3 software or with Leica thunder imaging system with Leica LAS-X software. Fluorescence was detected with mCherry, Texas Red (for dsRed labelled cells), and green fluorescent protein (GFP) filters.

### Image Analysis

All images were analyzed using FIJI software^42^.

### Quantification and statistical analysis

Statistical analyses were performed by the GraphPad Prism software (Prism 9). All experiments with only two groups and one dependent variable were compared using an unpaired t-test with Welch’s correction. Statistical data show mean ± S.E.M. Each dot plot value represents an independent embryo, and every experiment was conducted three times independently. The corrected total cell fluorescence (CTCF) was calculated using the formula ‘Integrated density whole – (area whole embryo x mean fluorescence background)’. This formula is loosely based on a method described for calculating cell-fluorescence^46^. To calculate the CTCF percentage for DsRed+ cells, the CTCF value of DsRed was divided by the total summation of CTCF values for both DsRed and GFP in each larvae. The same was done for calculating CTCF percentage for GFP+ cells. For fluorescence intensity quantification, the same parameters (excitation/emission, gain for detectors, lasers intensity) were applied for images and fluorescence intensity was measured using FIJI. The fold induction or the GFP-fold induction was calculated by dividing the average of the fluorescence intensity values in the light-exposed group by the average values of the non-exposed ones.

## Acknowledgments

The authors would like to acknowledge the Cell and Tissue Imaging Platform – PICT-IBiSA (member of France–Bioimaging – ANR-10-INBS-04) of the U934/UMR3215 of Institut Curie for help with light microscopy, in particular to Olivier Leroy for his help for achieving region-specific photoactivation. We also thank the members of the animal facility of Institut Curie for zebrafish care. We also thank Yohanns Bellaiche and all the members of the Hernandez lab for thoughtful and valuable discussions. We are grateful to Zilong Wen for providing all the immune and blood lineage switch lines used in this study. We are also thankful to Filipo Del Bene for providing the Hulk line. This work was supported by the Institut Curie, INSERM, CNRS, the Ville de Paris emergence program (2020 DAE 78), FRM amorçage (AJE201905008718), ATIP-Avenir starting grant R21045DS, ERC-StG Cytok-Gut 101041422, and the Laboratoire d’Excellence (Labex) DEEP (ANR-11-LBX-0044, ANR-10-IDEX-0001-02 PSL). R.E. was supported by the Springboard postdoctoral fellowship form Institut Curie and Labex DEEP.

## Author contribution

Conceptualization: R.E. and P.P.H.; Methodology and data collection: R.E., A.M., P.D. and G.G.; Writing - Original Draft: R.E.; Writing - Review & Editing: R.E. and P.P.H.; Funding Acquisition: P.P.H.

## Inclusion and diversity

We support inclusive, diverse and equitable conduct of research.

---

**Supplementary Figure 1/S1 related to figure 1:**
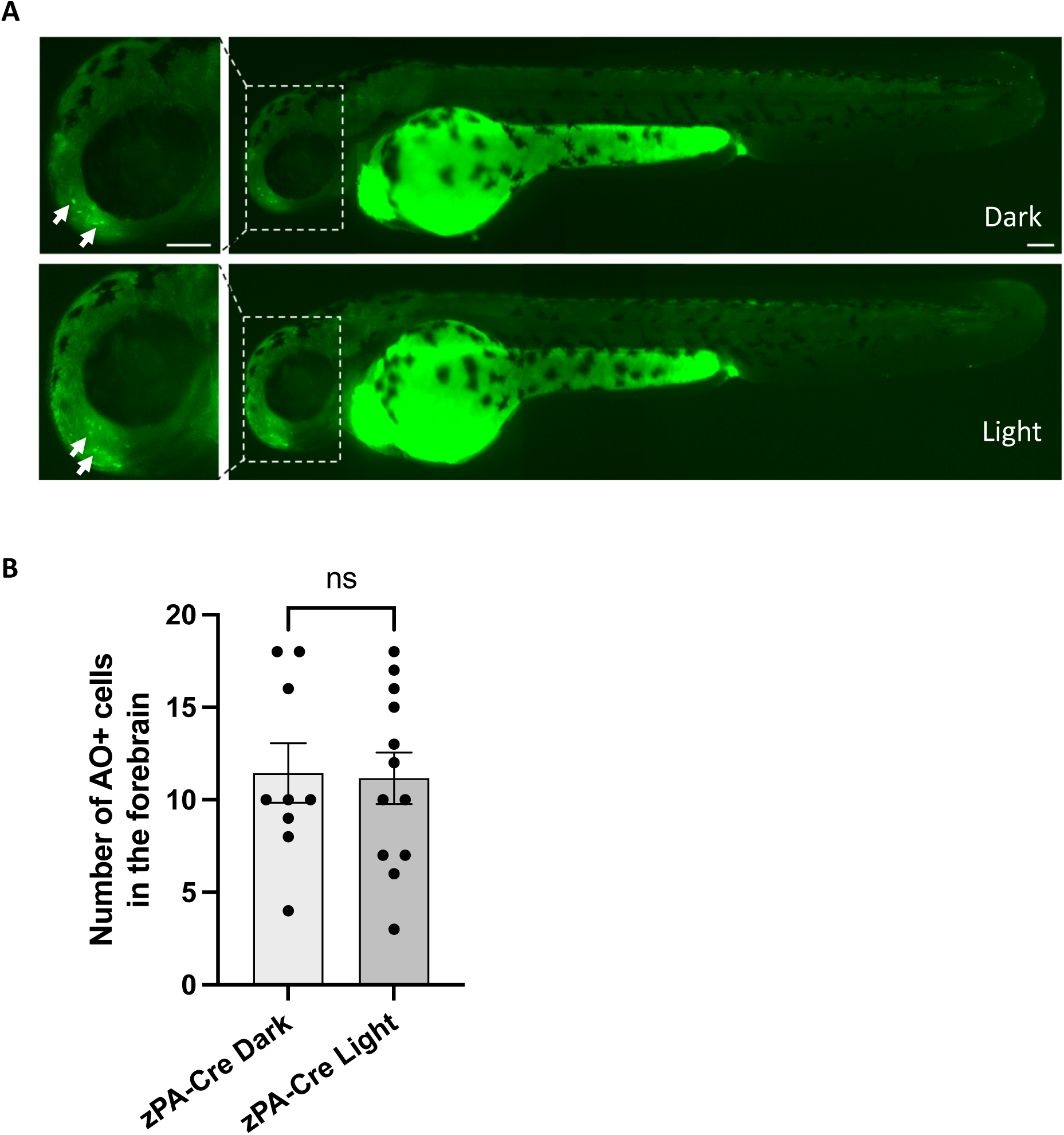
Assessing phototoxicity using Acridine Orange labeling. (A) Acridine orange (AO) staining is shown at 48 hpf in *Tg(ubb:PA-Cre)* embryos kept in the dark or exposed to blue light illumination at 6 hpf for 12 hours. Arrows highlight AO+ cells in the forebrain in closeups (left panel). Scale bar: 100 µm. (B) Quantification of the number of AO+ cells in the forebrain at 48 hpf of *Tg(ubb:PA-Cre)* embryos kept in the dark (n=9) or exposed to blue light illumination at 6 hpf for 12 hours (n=12). Mean ± SEM of the AO+ cells are shown. An unpaired t-test with Welch’s correction was used for this analysis. Ns, non-significant.

**Supplementary Figure 2/S2 related to figure 1:**
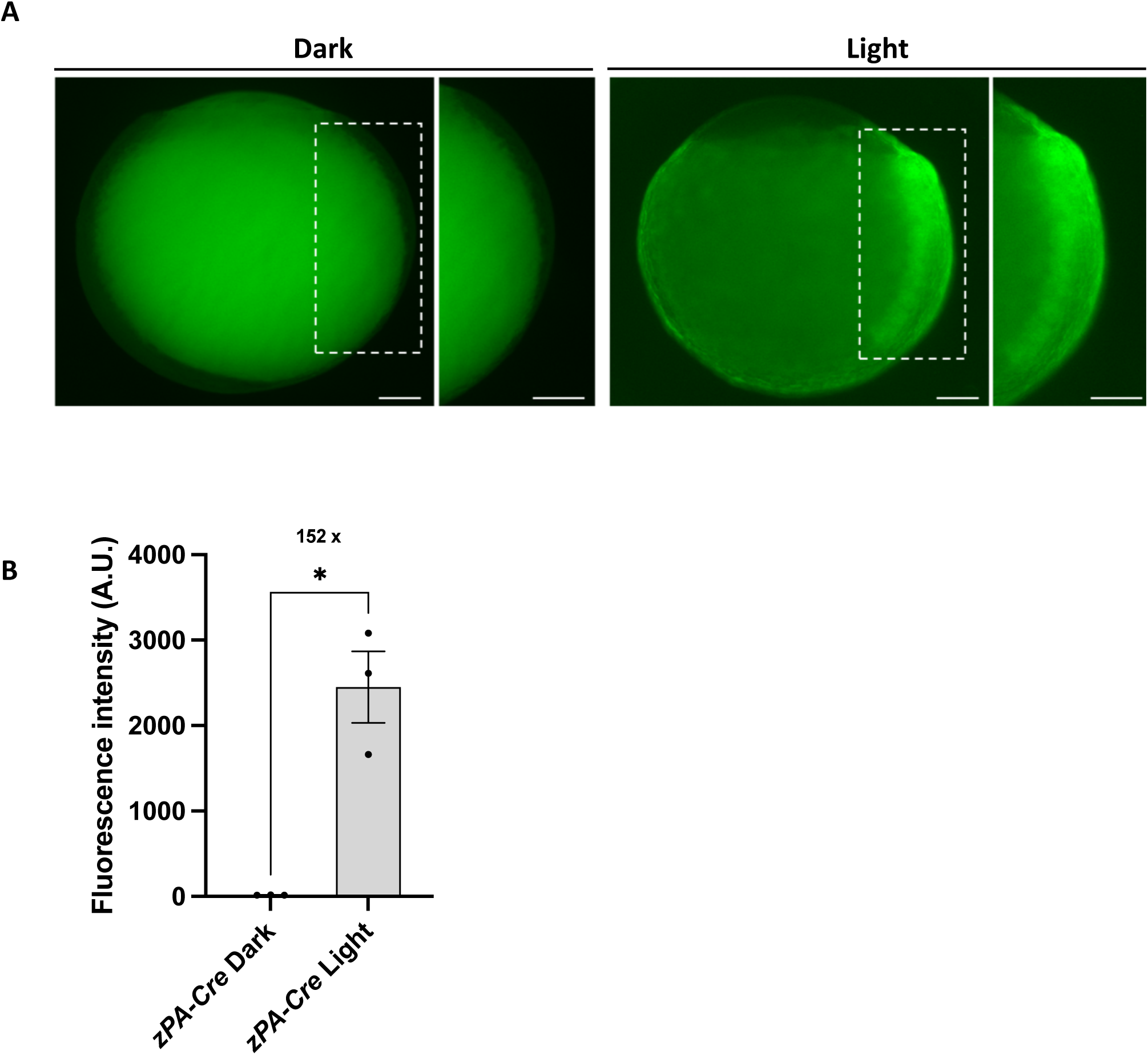
Assessing how fast GFP expression appears after photoactivation. (A) Fluorescent images at 8 hpf of *Tg(ubb:PA-Cre; ubb:Hulk)* embryos kept in the dark or exposed to blue light illumination. White dashes indicate the regions with closeups. Scale bar, 100 µm. (B) Quantification of fluorescence intensity at 8 hpf of *Tg(ubb:PA-Cre; ubb:Hulk)* embryos kept in the dark (n=3) or exposed to blue light illumination at the 1-cell stage (n=3). 152 x represents the fold induction comparison between the two groups. Mean ± SEM of the fluorescence intensity is shown. Unpaired t-test with Welch’s correction was used for this analysis. *p ≤ 0.1. A.U.; Arbitrary Unit.

**Supplementary Figure 3/S3:**
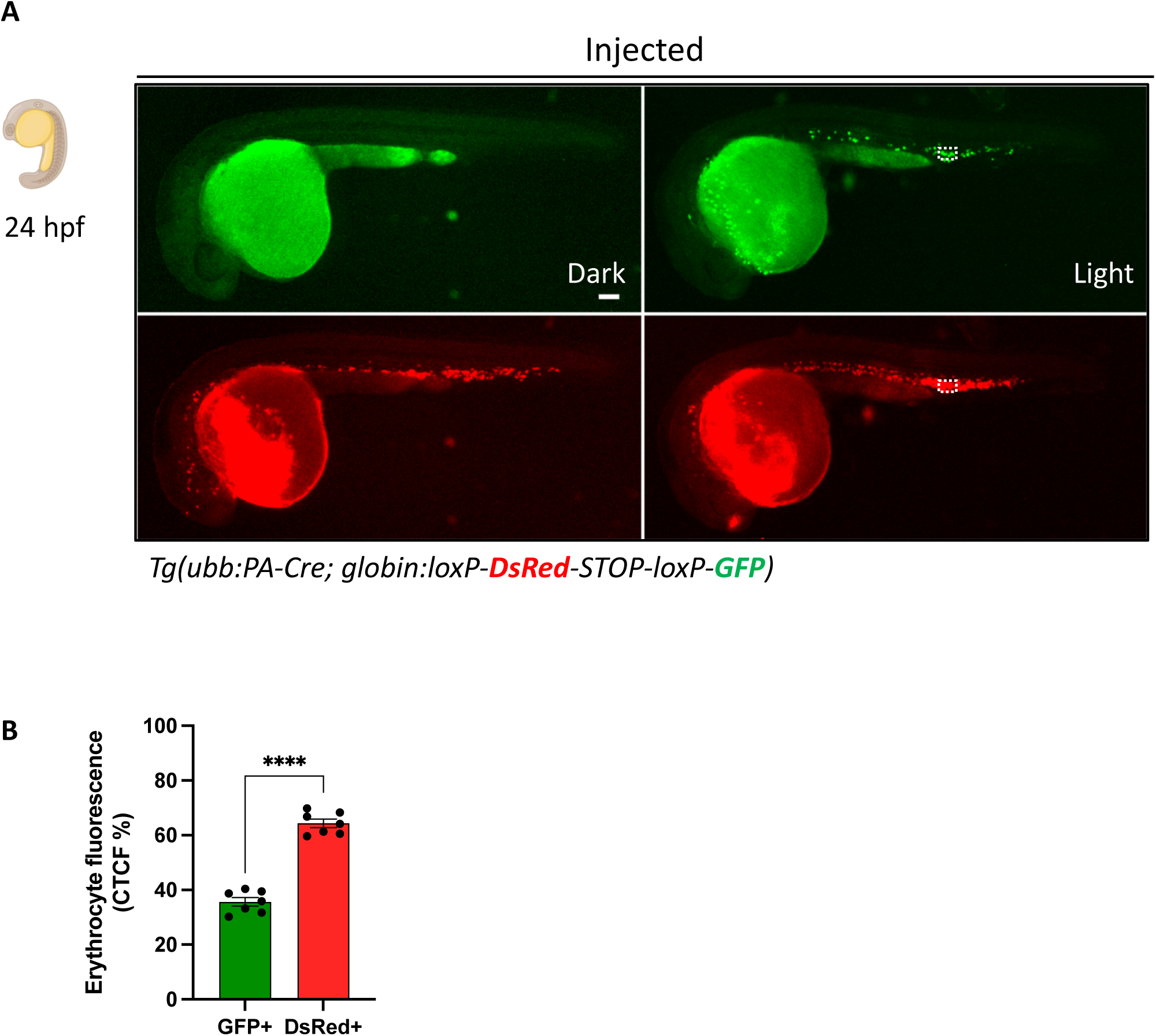
Testing the functionality of zPA-Cre under the kdrl promoter. (A) Representative fluorescent images at 24 hpf of *Tg(globin:Switch)* embryos injected with the kdrl-PA:Cre construct at one-cell stage and kept in the dark (left panel) or exposed to blue light illumination at 6 hpf for 12 hours (right panel). White dashes represent the area of the CTCF quantifications. Scale bar: 100 µm. (B) Quantification of erythrocyte fluorescence intensity was measured at 24 hpf in photoactivated *Tg(ubb:PA-Cre; globin:Switch)* embryos (n=7). Mean ± SEM of the DsRed+ and GFP+ corrected total cell fluorescence (CTCF) percentage is shown. Unpaired t-test with Welch’s correction was used for this analysis. ***p ≤ 0.0001.

**Supplementary Figure 4/S4 related to figure 4:**
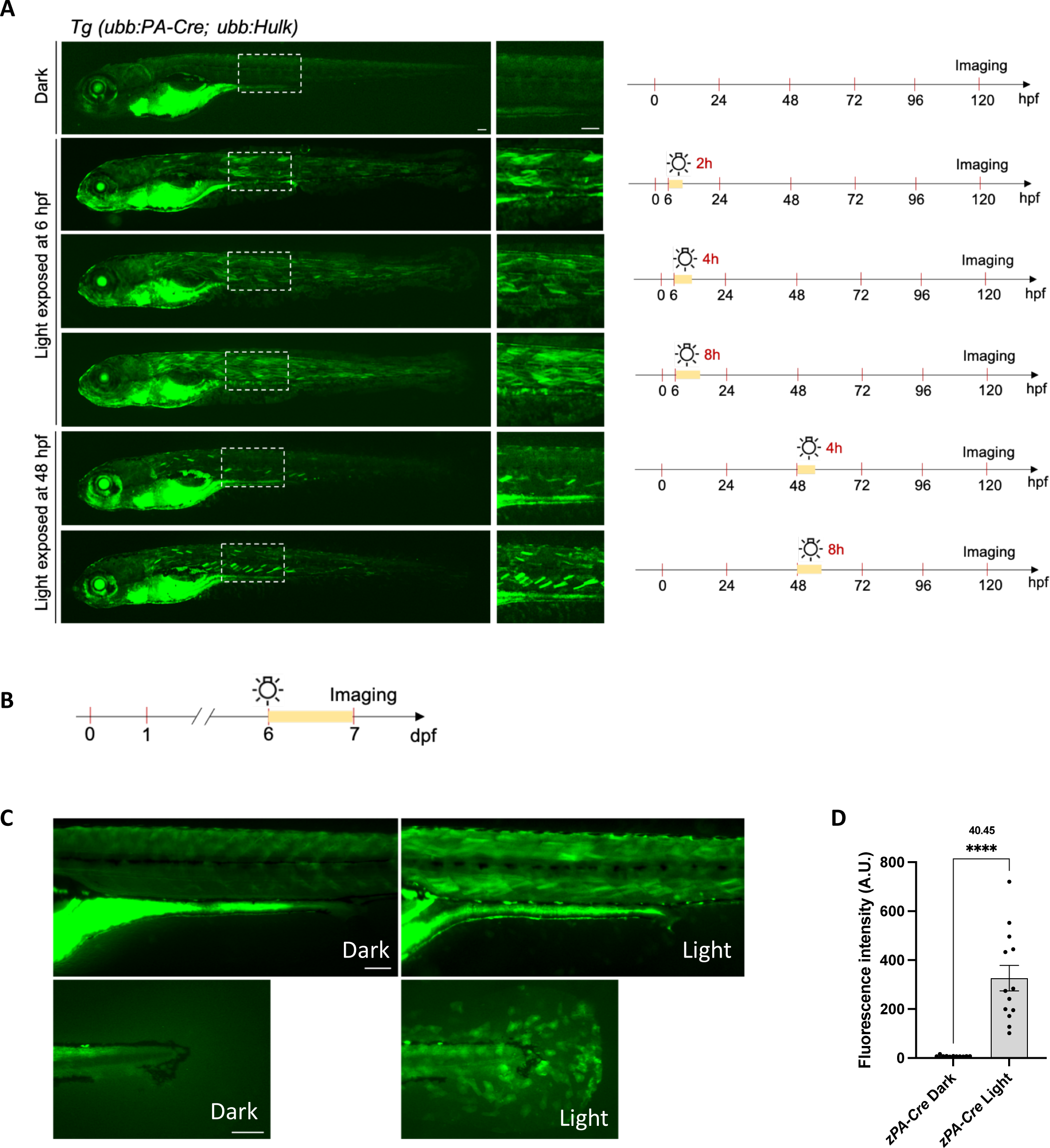
Assessing the photoactivation of the zPA-Cre system in response to different light doses and beyond embryonic stages. (A) Representative fluorescent images at 120 hpf of *Tg (ubb:PA-Cre; ubb:Hulk)* embryos kept in the dark or exposed to blue light illumination at 6 hpf for 2 hours, 4 hours, or 8 hours, and at 48 hpf for 4 hours and 8 hours (left panel). White dashes indicate the regions with closeups at the right. Scale bar, 100 µm. Schematic representation of the strategy used to induce genetic recombination in zebrafish *Tg(ubb:PA-Cre; ubb:Hulk)* in response to different light doses (right panel). (B) Schematic representation of the strategy used to induce genetic recombination in *Tg(ubb:PA-Cre; ubb:Hulk)* zebrafish at the larval stage. (C) Representative fluorescent images at 7 dpf of *Tg (ubb:PA-Cre; ubb:Hulk)* larvae kept in the dark or exposed to blue light illumination at 6 dpf overnight. The upper panel shows a part of the larvae’s trunk, and the bottom panel shows the tail. Scale bar, 100 µm. (D) Quantification of fluorescence intensity at 7 dpf of *Tg (ubb:PA-Cre; ubb:Hulk)* larvae kept in the dark (n=11) or exposed to blue light illumination at 6 dpf overnight (n=13). 40.45 x represents the fold induction comparison between the two groups. Mean ± SEM of the fluorescence intensity is shown. An unpaired t-test with Welch’s correction was used for this analysis. ****p ≤ 0.0001. A.U.; Arbitrary Unit.

**Supplementary Video 1 related to Figure 2:**
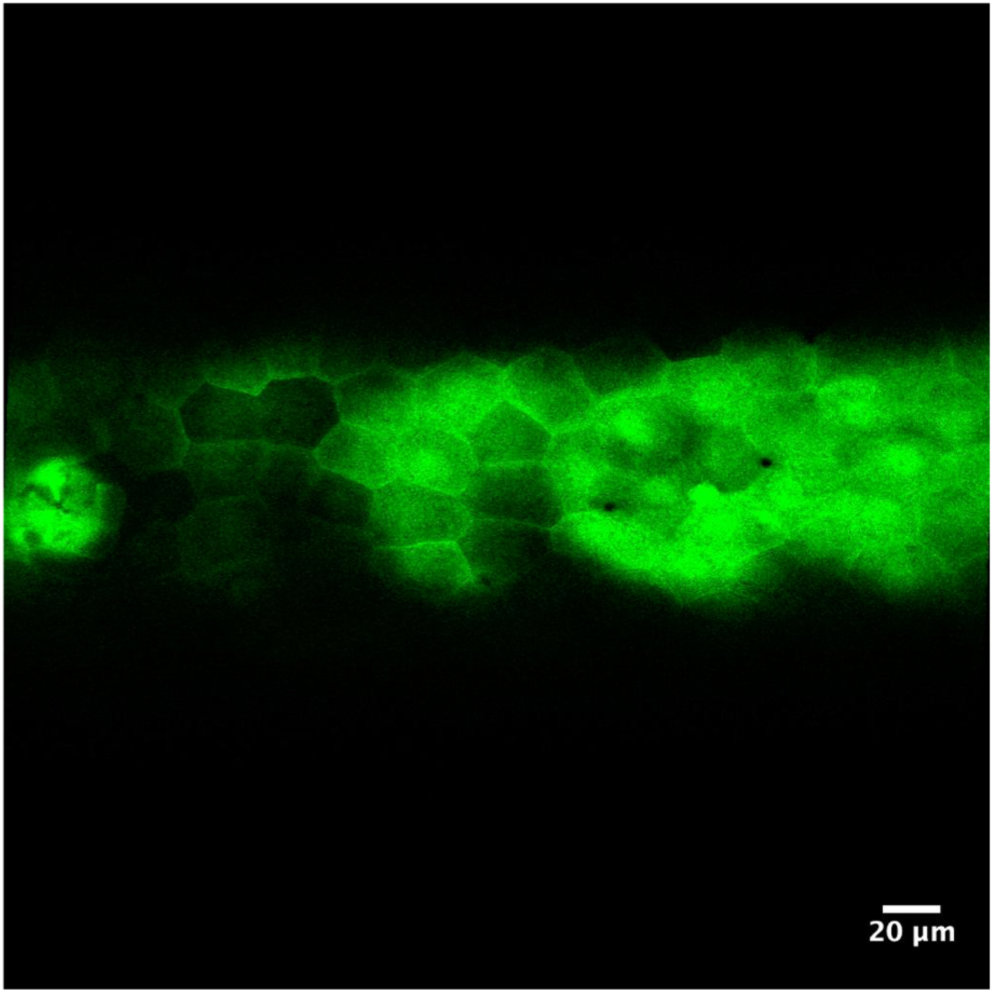
Assessing the light-induced genetic recombination efficiency in multiple tissues in a light exposed zebrafish embryo. Fluorescence imaging of photoactivated *Tg(ubb:PA-Cre; ubb:Hulk)* at 72 hpf. The video is assembled from different Z-stack projections. Lateral view; anterior to the left and dorsal to the top. Scale bar: 20 µm.

**Supplementary Videos 2 related to Figure 6:**
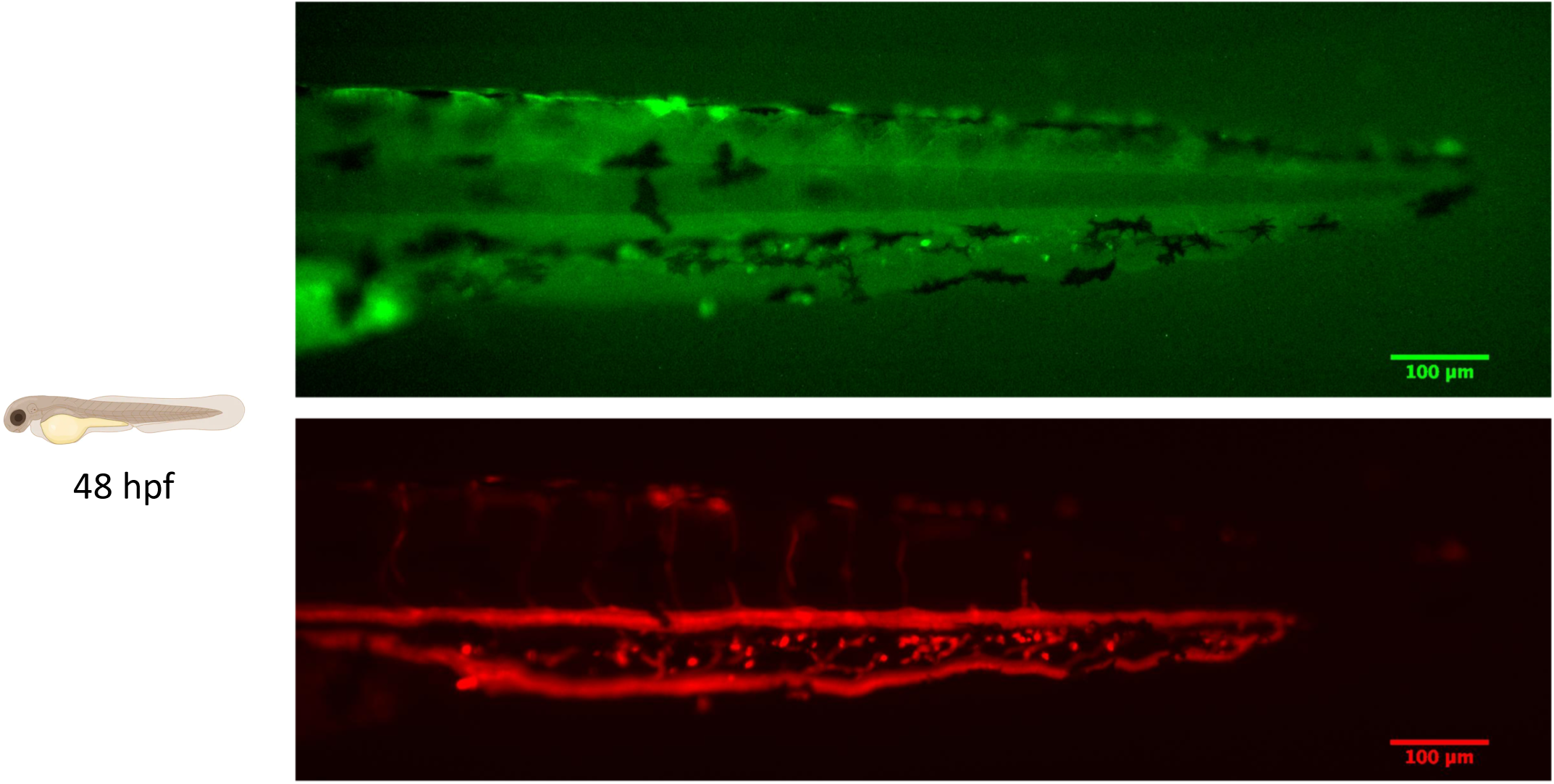
Testing the the spatio-temporal control efficiency of the zPA-Cre system using blue laser illumination. Real-time imaging in *Tg(ubb:PA-Cre; globin:Switch)* at 48 hpf. A 22 hpf embryo was used to induce local photoactivation by a 488 nm laser for 5 minutes within a 25 µm diameter circle in the posterior blood island (PBI). Lateral view; anterior to the left and dorsal to the top. Scale bar: 100 µm.

**Supplementary Videos 3 related to Figure 6:**
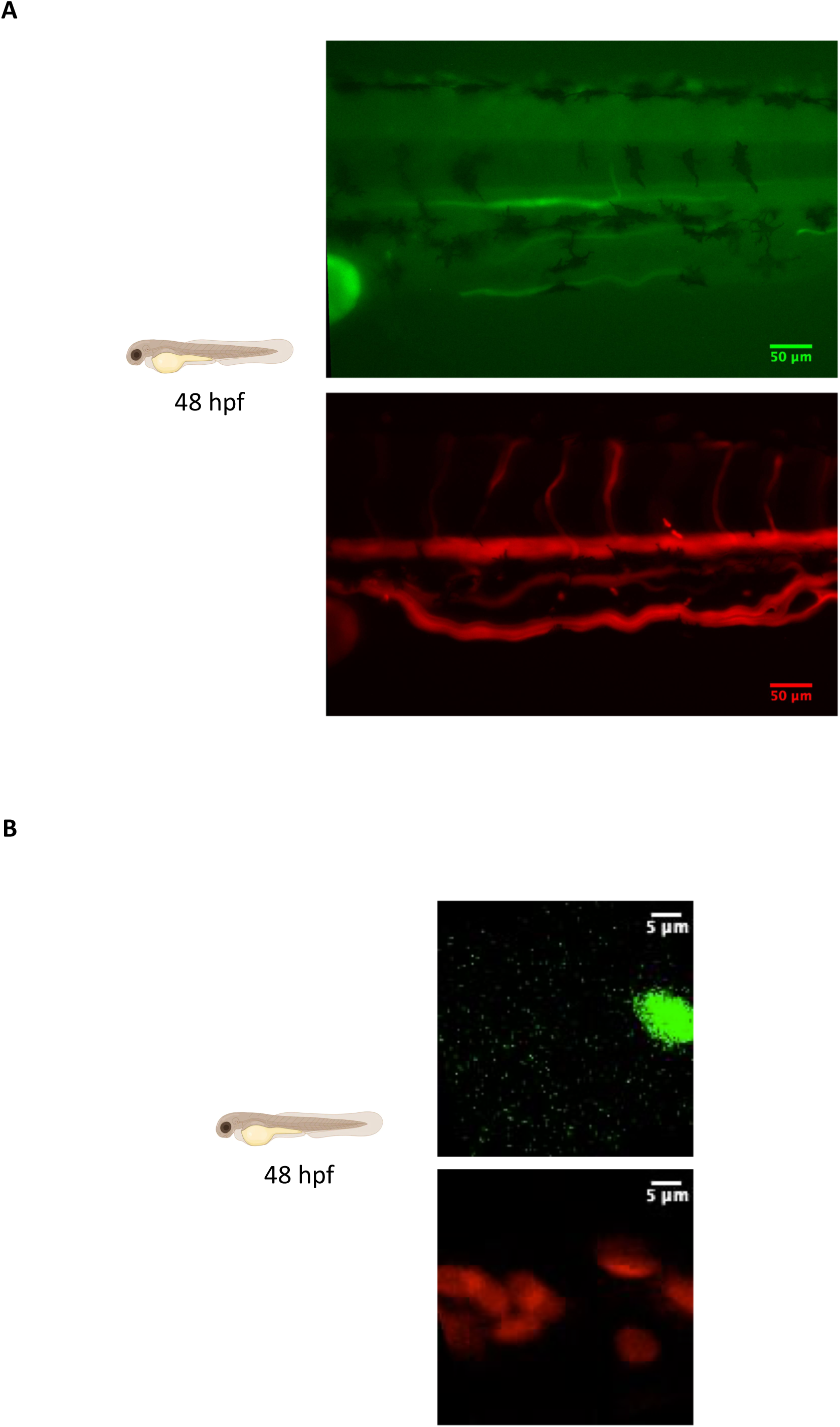
Assessing the single cell targeting efficiency of the zPA-Cre system with using infrared laser illumination. (A) Real-time imaging in *Tg(ubb:PA-Cre; globin:Switch)* at 48 hpf. A 22 hpf embryo was used to induce photoactivation by a 720 nm two-photon laser for 45 seconds in an 8x8 µm single erythrocyte in the posterior blood island (PBI). Lateral view; anterior to the left and dorsal to the top. Scale bar: 50 µm. Time-lapses of higher magnification are shown in (B). (B) Higher magnification of real-time imaging in *Tg(ubb:PA-Cre; globin:Switch)* at 48 hpf. A 22 hpf embryo was used to induce photoactivation by a 720 nm two-photon laser for 45 seconds in an 8x8 µm single erythrocyte in the posterior blood island (PBI). Lateral view; anterior to the left and dorsal to the top. Scale bar: 5 µm.

**Supplementary Note.**
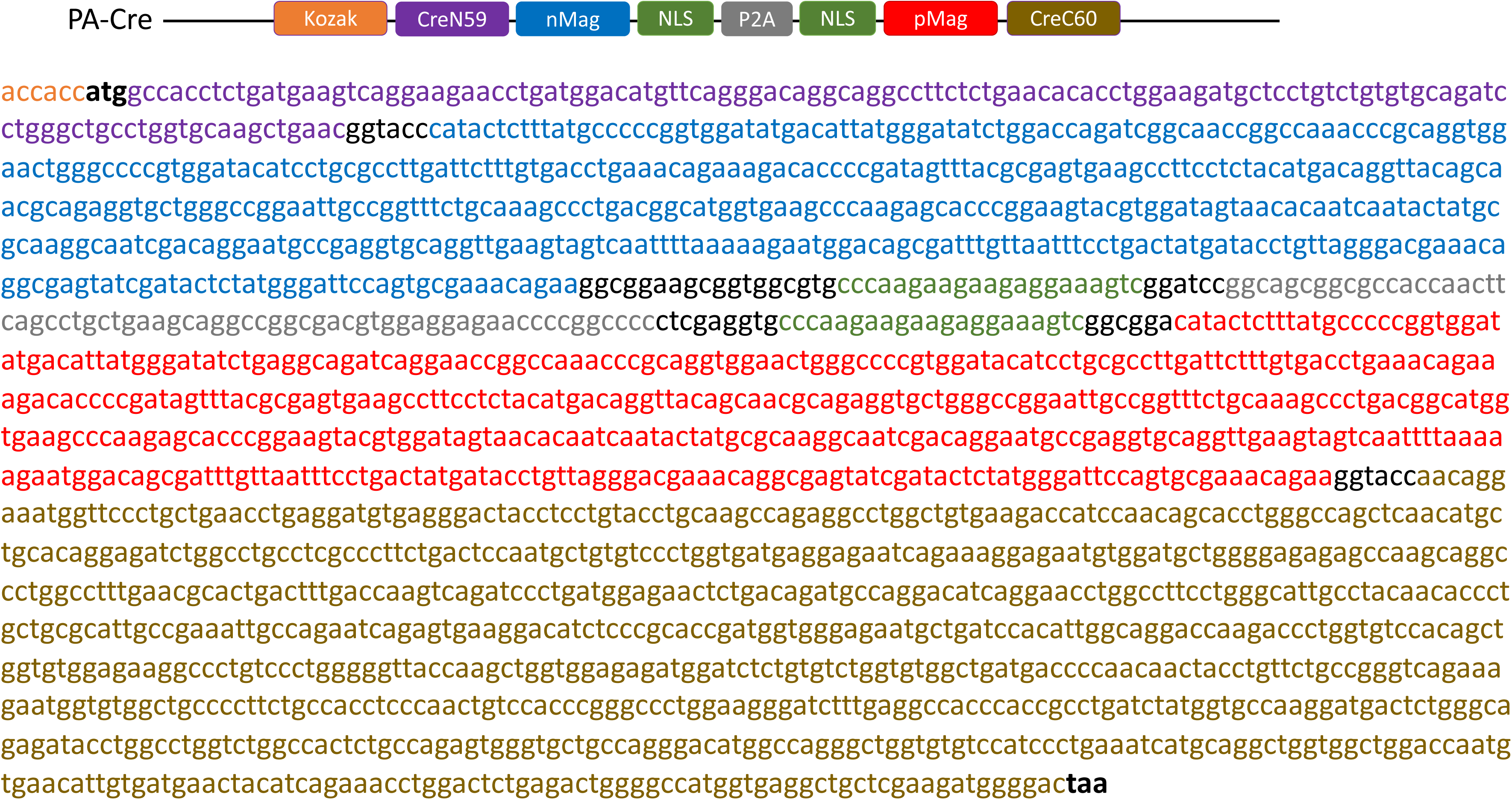
Nucleic acid sequences of PA-Cre used in the this study.

